# Proteomics analysis of autophagy cargos reveals distinct adaptations in PINK1 and LRRK2 models of Parkinson disease

**DOI:** 10.1101/2022.10.03.510717

**Authors:** Juliet Goldsmith, Alban Ordureau, C Alexander Boecker CA, Madeleine Arany, J Wade Harper, Erika LF Holzbaur

**Affiliations:** Department of Physiology, University of Pennsylvania Perelman School of Medicine, Philadelphia PA 19104 USA; Department of Cell Biology, Blavatnik Institute, Harvard Medical School, Boston, MA, 02115 USA; Cell Biology Program, Sloan Kettering Institute, Memorial Sloan Kettering Cancer Center, New York, NY, 10065 USA; Department of Neurology, University Medical Center Goettingen, 37077 Goettingen, Germany; Aligning Science Across Parkinson’s (ASAP) Collaborative Research Network, Chevy Chase, MD, 20815 USA

**Keywords:** α-synuclein, extracellular vesicles, LRRK2, mitophagy, Parkinson disease, PINK1, secretory autophagy

## Abstract

Autophagy is essential for neuronal homeostasis, while defects in autophagy are implicated in Parkinson disease (PD), a prevalent and progressive neurodegenerative disorder. We used unbiased proteomics to compare cargos degraded by basal autophagy in the brain from two mouse models of PD, PINK1^-/-^ and LRRK2^G2019S^ mice. We find evidence for the upregulation of adaptive pathways to support homeostasis in both PD models. In PINK1^-/-^ mice, we observed increased expression of the selective receptor BNIP3 along with evidence of engagement of other alternative pathways for mitophagy. Despite these changes, we find the rate of autophagic flux in PINK1^-/-^ neurons is decreased. In LRRK2^G2019S^ mice, hyperactive kinase activity known to impair autophagosomal and lysosomal function results in increased secretion of extracellular vesicles and autophagy cargo. In support of this observation, we find reduced levels of PIKFYVE, a negative regulator of extracellular vesicle secretion, in both brain and cortical neurons from LRRK2^G2019S^ mice. Thus, distinct adaptive pathways are activated to compensate for perturbations induced by either loss of PINK1 or hyperactivation of LRRK2. Our findings highlight the engagement of compensatory pathways to maintain homeostasis in the brain, and provide insights into the vulnerabilities these compensatory changes may introduce that may further contribute to PD progression.

## INTRODUCTION

Autophagy is an evolutionarily conserved process for the clearance and recycling of proteins and organelles. Neurons have a unique dependence on autophagy, as impairment of this pathway leads to either neurodevelopmental or neurodegenerative phenotypes [1–3]. Cellular, genetic, and pathological data point to two major categories of autophagy in neurons: selective autophagy and macroautophagy. Acute stress induces the selective autophagic targeting of dysfunctional organelles or aggregated proteins for turnover via regulated pathways: for example, mitophagy, lysophagy and aggrephagy for the clearance of damaged mitochondria, lysosomes or aggregated proteins respectively [4]. These pathways generally involve damage-sensing, leading to induction of a downstream response. For example, mitochondrial damage leads to the stabilization of the kinase PINK1 on the outer mitochondrial membrane (OMM), which in turn triggers a feedforward activation of the E3 ubiquitin ligase Parkin, leading to widespread ubiquitination of mitochondrial proteins and recruitment of ubiquitin-binding receptors such as OPTN that serve as a platform leading to the engulfment of the damaged organelle by a double-membrane autophagosome [5–8].

In addition to these stress-induced mechanisms, there is robust cellular and in vivo evidence for a requirement of basal macroautophagy in neurons [9,10]. We previously used proteomics to identify the major cargos engulfed by autophagic vesicles in the brain under basal conditions. Western blotting and live imaging studies of induced human neurons and primary rodent neurons confirmed that mitochondrial fragments enriched in nucleoids and synapse-related proteins were highly enriched in autophagic vesicles formed within neurons under basal conditions [11]. Both in vitro and in vivo, autophagosomes form constitutively at synaptic sites and the axon terminus where they engulf their mitochondrial and synaptic protein cargos, and then undergo a stereotypical pattern of highly regulated transport toward the soma, driven by the molecular motor cytoplasmic dynein [12–15]. Autophagosome transport is critical for maturation, which is necessary for the efficient degradation of engulfed cargos [16,17].

Parkinson disease (PD) is a progressive neurodegenerative disease primarily affecting dopaminergic neurons in the substantia nigra pars compacta. It presents as a movement disorder in patients with an average age of onset at 60. Disrupted autophagy is strongly associated with PD. Histopathologically, abnormal autophagosomes are accumulated in post-mortem patient brain tissue independent of the etiology [18]. Functional autophagy is responsible for clearing α-synuclein aggregates that can form the characteristic pathological Lewy bodies found in PD, suggesting that the defects in autophagy may precede Lewy body accumulation [19,20].

Genetic evidence also implicates disruption of autophagy in the progression of PD. Roughly 15% of PD cases are familial, with mutations in the leucine-rich repeat kinase 2 (*LRRK2*) gene resulting in hyperactive LRRK2 kinase activity being one of the most common causes. Additionally, there is evidence of elevated LRRK2 kinase activity in idiopathic PD cases [21]. Hyperactive LRRK2 leads to increased phosphorylation of RAB proteins, direct LRRK2 kinase targets [22,23] and key regulators of vesicle trafficking pathways [24]. Elevated LRRK2 activity in neurons is sufficient to disrupt both the transport and the maturation of autophagic vesicles in the axon, leading to a block in autophagic degradation [25,26]. Rare familial mutations in the machinery required for the selective autophagic degradation of damaged mitochondria (mitophagy), including mutations in PINK1, cause more acute forms of PD with earlier age of onset and higher penetrance than the LRRK2 mutations. While the genetic evidence suggests that impairments to the autophagy pathway are causative for disease, the late age of onset in PD patients implicates the involvement of additional disease drivers, modifiers, or compensatory mechanisms that lead to the slow accumulation of damage over time.

Here, we report changes in autophagy cargo observed in two different models of PD. We used unbiased proteomics to analyze autophagic cargos in *PINK1* knock-out mice and *LRRK2*^G2019S^ knock-in mice, which model one of the most severe phenotypes of PD and one of the most common familial mutations in PD, respectively. In both models, we find evidence for the upregulation of compensatory pathways for organelle quality control and protein clearance. Specifically, loss of PINK1 in mice leads to compensatory changes in mitophagy machinery such as the upregulation of BNIP3. We find that mitochondria are still engulfed within autophagic vesicles (AVs) and mitochondria can still be cleared following damage in the absence of PINK1, although autophagosome degradation occurs less efficiently. In LRRK2^G2019S^ mice we find evidence of increased secretion, which we hypothesize supports neuronal homeostasis by removing damaged organelles or aggregated proteins from the affected neurons. Overall, we highlight a common theme of engagement of compensatory pathways in response to PD-associated mutations in order to maintain organelle quality control and proteostasis in neurons. The engagement of compensatory mechanisms may explain the limited phenotypes observed in mouse models but may also introduce potentially injurious downstream consequences. Based on our observations in these mouse models, we propose that similar compensatory changes may contribute to age-dependent onset of disease in patients carrying pathogenic mutations in PINK1 or LRRK2.

## RESULTS

### Unbiased proteomics reveal changes to autophagic cargo in the brain of PINK1^-/-^ mice

The accumulation of PINK1 on the outer mitochondrial membrane of damaged mitochondria is an important step in activating Parkin-dependent mitophagy. However previous studies in Drosophila and mouse indicate that under basal conditions, PINK1 and/or Parkin contribute minimally to mitochondrial turnover [10,11,27]. Therefore, we were curious to define the impacts of PINK1 loss on mitochondrial turnover by autophagy under basal conditions, and whether the loss of PINK1 *in vivo* has other appreciable effects on mitophagy or autophagy more broadly.

We performed autophagic vesicle enrichment using differential centrifugation from whole brains of control (CJ n=5), or PINK1^-/-^ (n=9) mice. Following isolation, the autophagic vesicle (AV) fraction was divided, with half treated with a Proteinase-K digestion step to degrade proteins associated with the external membrane of AVs. This step enriches for proteins that are protected by the double membrane of the autophagic vesicle and thus remain undigested by Proteinase-K (Fig. 1A). We previously characterized the specificity and selectivity of this approach [11].

**Figure 1.**
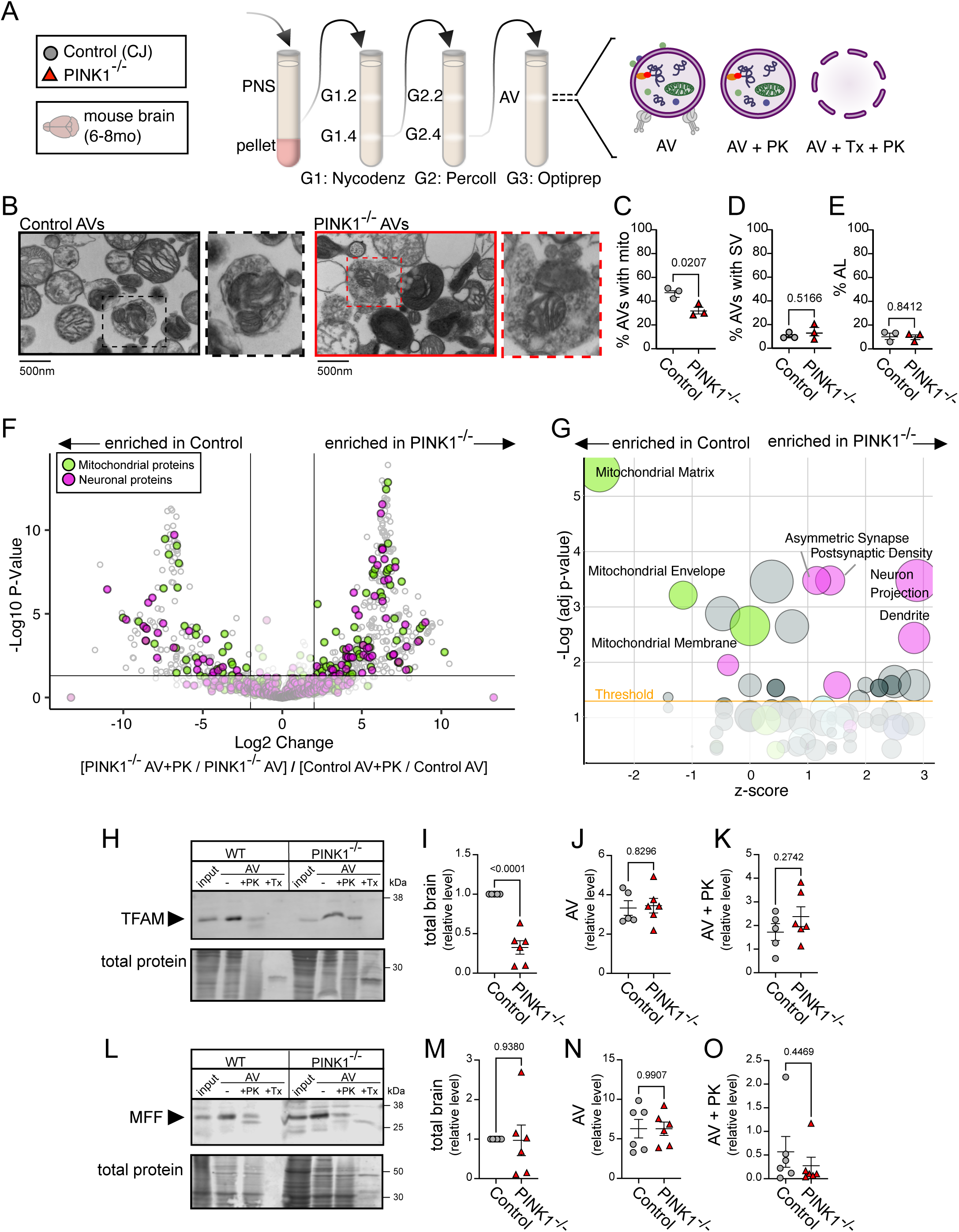
Characterization of PINK1^-/-^ brain-derived autophagosome cargo indicates mitochondria remain basal autophagy cargo **A.** Schematic representing the differential ultracentrifugation protocol and Proteinase-K digestion step for the enrichment of autophagic vesicles from control and PINK1^-/-^ mouse brain following euthanasia at 6-8 months of age. **B.** Representative electron micrographs of AVs derived from WT and PINK1-/- mouse brains. Insert highlights an identified AV containing a mitochondrial fragment and synaptic vesicle like structures. **C-E.** Quantification of AVs containing (**C**) mitochondria, (**D**) synaptic vesicle like structures, or (**E**) electron dense autolysosomes from EM images (Control n = 3; PINK1^-/-^ n = 3, >400 events counter per biological replicate). **F.** Volcano plot analysis of the ratio of AV+PK to AV fractions derived from the brain of PINK1^-/-^ relative to control mice. Black lines indicate significance thresholds of a p-value < 0.05 and a Log2 fold change > 2, and points are colored magenta if representing a neuronal or synapse associate protein, and green if representing a mitochondrial protein. Control n = 5, PINK1^-/-^ n = 9. **G.** Bubble plot describing the significance and directionality of change of Gene Ontology terms from proteins whose cargo score was significantly changed between PINK1^-/-^ and control. Each bubble represents a unique GO term, colored magenta if representing a neuronal or synapse associate term, and green if representing a mitochondrial term. A significance threshold of an adjusted p-value <0.05 is marked in by a yellow line. The z-score represents the overall directionality of the cargo score, with a score of 0 indicating equal enrichment in both control and PINK1^-/-^. The size of the circle corresponds to the number of proteins identified in the GO term. **H-K.** Representative immunoblot (**H**) and quantifications (mean ± SEM, unpaired t test) of the levels of TFAM in control and PINK1^-/-^ (**I**) total brain, (**J**) AV fraction, and (**K**) AV+PK fraction lysate, normalized to total protein and control total brain levels. Control n = 5, PINK1^-/-^ n = 6. **L-O.** Representative immunoblot (**L**) and quantifications (mean ± SEM, unpaired t test) of the levels of MFF in control and PINK1^-/-^ (**M**) total brain, (**N**) AV fraction, and (**O**) AV+PK fraction lysate, normalized to total protein and control total brain levels. Control n = 6, PINK1^-/-^ n = 6.

Electron microscopy (EM) of enriched AV fractions from both PINK1^-/-^ and control mice revealed double membrane vesicles surrounding mitochondria or synaptic vesicle-like structures, as well as electron-dense vesicles we designated as autolysosomes (AL) (Fig. 1B). The percentages of each category found in AV fractions from control mice were consistent with our previously published findings [11]. A significantly smaller proportion of brain-derived AVs from PINK1^-/-^ animals contained mitochondria (Figure 1C), while the percentage of AVs containing synaptic vesicle-like structures and the percentage of autolysosomes remained unchanged (Fig. 1D,E).

Proteomic analysis was performed on the AV fraction and the Proteinase-K treated (PK) fraction from mice of both genotypes (Tables S1, S2). We detected increased numbers of peptides in both the total and PK-treated fractions isolated from PINK1^-/-^ brain as compared to parallel control samples from wild type mice (Fig. S1A, B). We noted that 1704 proteins were significantly higher in the PINK1^-/-^ AV fraction compared to only 6 proteins significantly higher in AVs isolated from control mice (Fig. S1C). Similarly, 1644 proteins were significantly higher in the PK-treated AV fraction from PINK1^-/-^ mice compared to only 7 proteins significantly higher in the control fraction (Fig. S1D). Mitochondria-associated terms were significantly upregulated in both the PINK1^-/-^ AV and PK fractions by gene ontology (GO) analysis (Fig. S1E,F).

The large differences in peptide abundances between AVs isolated from control and PINK1^-/-^ masks changes to cargo selection. Because we performed proteomic analysis on both the AV and PK fractions, we are able to internally normalize for each independent enrichment preparation. We compared the cargo scores, defined as the ratio of peptide abundance in the PK fraction to the total AV fraction [11], from PINK1^-/-^ mice to control mice to compare whether the composition of autophagic cargos is altered due to the loss of PINK1 (Table S3). We highlighted mitochondrial proteins and neuronal or synapse associated proteins (Fig. 1F). While individual mitochondrial or synapse associated proteins were found at either increased or decreased levels in PINK1^-/-^ derived AVs, GO analysis highlights the overall observation that neuronal terms are more enriched in PINK1^-/-^ derived AVs, while mitochondrial terms are less enriched (Fig. 1G). These findings are consistent with EM quantification, which revealed proportionally fewer autophagic vesicles containing mitochondria-like structures.

Mitochondria that contain TFAM-positive nucleoids are constitutively engulfed by autophagosomes in the absence of mitochondrial damage [11], so we investigated how the loss of PINK1 may affect TFAM clearance. We ran a fraction of total brain sample, and equally loaded volumes of the AV, AV+PK, and the negative control triton-X and PK treated AV fractions on SDS-PAGE gel and immunoblotted for our cargo of interest. The total protein levels observed are reflective of the efficacy of the PK treatment. There was a decrease in TFAM levels in whole brain samples from PINK1^-/-^ mice, but the amount of TFAM found in the AV and PK fractions was unchanged (Fig. 1H-K). Similarly, levels of MFF, another mitochondrial protein turned over by constitutive autophagy, were consistent across autophagosomes from control and PINK1 mice (Fig. 1L-O). This suggests that the basal engulfment of nucleoid-enriched mitochondrial fragments that occurs in the distal axon in the absence of damage is unaffected by loss of PINK1^-/-^.

### PINK1 loss results in increased levels of selective mitophagy receptors within AVs

We looked for evidence of compensatory changes to the mitophagy pathway, asking whether known receptors for selective autophagy or mitophagy were found at altered levels within PINK1^-/-^ derived AVs. Four mitophagy-associated proteins — BNIP3, FUNDC1, FUNDC2, and MFN1 — were found to have four-fold or greater cargo scores and p-values lower than 0.05 (Fig. 2A). Other autophagy-associated proteins, including WDFY3 and TOLLIP, were also increased in both the total AV fraction and PK-treated fractions from PINK1^-/-^ mice relative to controls (Fig. S2A, B).

**Figure 2.**
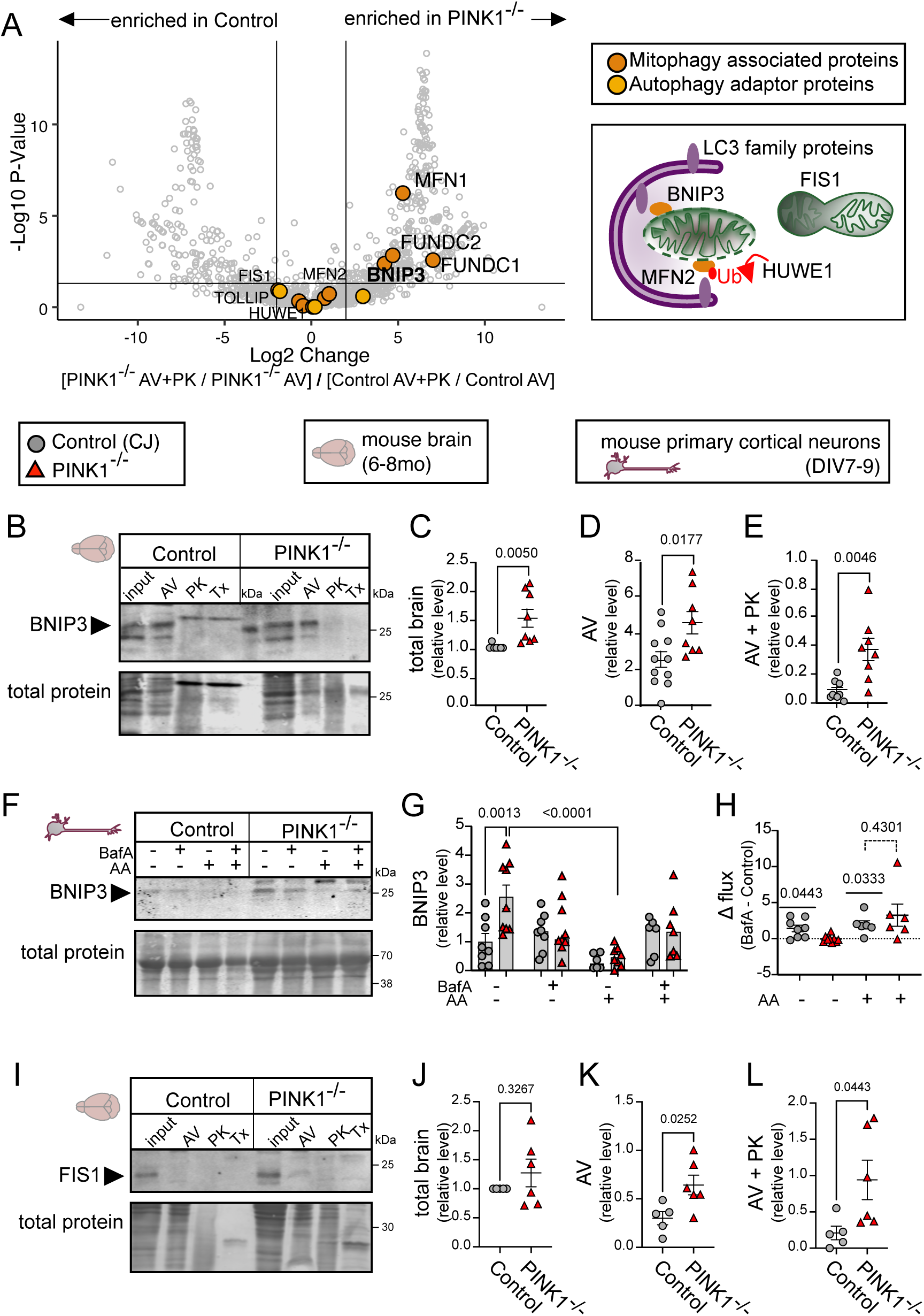
Alternate mitophagy receptors in brain and neurons are increased in the absence of PINK1 **A.** Identical volcano plot analysis as in Figure 1F comparing the change in cargo score from PINK1^-/-^ relative to control brain derived AVs, with mitophagy associated proteins and autophagy receptor proteins highlighted in orange and gold respectively. A cartoon schematic of alternate mitophagy pathways is diagrammed. **B-E.** Representative immunoblot (**B**) and quantifications (mean ± SEM, unpaired t test) of the levels of BNIP3 in control and PINK1^-/-^ (**C**) total brain, (**D**) AV fraction, and (**E**) AV+PK fraction lysate, normalized to total protein and control total brain levels. Control n = 11, PINK1^-/-^ n = 8. **F,G.** Representative immunoblot (**F**) and (**G**) quantification (mean ± SEM, two-way ANOVA with Šídák’s multiple comparisons test) of the levels of BNIP3 in control and PINK1^-/-^ primary neuron lysate from a mitophagy flux assay with Bafilomycin A (100 nM for 2 h) and Antimycin A (15 nM for 2 h), normalized to total protein and control untreated levels. Control n ≥ 6, PINK1^-/-^ n ≥ 7. **H.** Graph showing the change in flux levels of BNIP3 from the paired Bafilomycin A treatment and untreated control and PINK1^-/-^ primary neurons, where 0 indicates no change in protein levels. One sample t and Wilcoxon test indicated by the line above the data points compares the mean ± SEM to the theoretical mean of 0. Dashed line between conditions indicates one way ANOVA with Šídák’s multiple comparisons test. Control n ≥ 6, PINK1^-/-^ n ≥ 6. **I-L.** Representative immunoblot (**I**) and quantifications (mean ± SEM, unpaired t test) of the levels of FIS1 in control and PINK1^-/-^ (**J**) total brain, (**K**) AV fraction, and (**L**) AV+PK fraction lysate, normalized to total protein and control total brain levels. Control n = 5, PINK1^-/-^ n = 6.

We compared the levels of the mitophagy receptor protein BNIP3 in whole brain by immunoblot (Fig. 2B-E) and found significantly higher levels of this mitophagy receptor in total brain lysate from PINK1^-/-^ mice relative to control mice (Fig. 2C). We also found significantly increased levels of BNIP3 in both total AVs and PK-treated AVs from PINK1^-/-^ mice, suggesting that this mitophagy receptor is upregulated in order to compensate for the loss of PINK1 (Fig. 2D, E).

As immunoblotting indicated increased levels of BNIP3 in the whole brain of PINK1^-/-^ animals, we next investigated whether primary neurons from this model also displayed increased levels of BNIP3. PINK1^-/-^ primary cortical neurons had increased total levels of BNIP3 (Fig. 2F, G). To understand whether BNIP3 is engaged in targeting mitochondria to the autophagosome, we employed an autophagy flux assay. Briefly, a flux assay monitors the levels of proteins at baseline and whether they accumulate in response to the V-ATPase inhibitor Bafilomycin A, which blocks autophagosomal-lysosomal degradation. In the absence of mitochondrial damage, there was little turnover of BNIP. However, upon acute mitochondrial damage with Antimycin A treatment, we found that levels of BNIP3 decreased. This decrease was blocked by Bafilomycin A in both the control and the PINK1^-/-^ neurons (Fig. 2G, H), indicating that BNIP3 is turned over by the autophagolysosomal pathway. Therefore, as we see higher levels of BNIP3 expression in both PINK1^-/-^ brain lysates and in primary cortical neurons, as well as robust BNIP3 degradation by autophagy in neurons, these data suggest that this protein is functioning in a compensatory pathway for mitochondrial clearance in PINK1^-/-^ mice.

We next asked whether other mitophagy related proteins observed at increased levels in AVs from PINK1^-/-^ mice also were found at increased levels in the brain. HUWE1, an E3 ligase that regulates PINK1^-/-^ independent mitophagy by ubiquitylating MFN2 and recruiting AMBRA1 to initiate autophagosome formation [28], was significantly increased in the PINK1^-/-^ AV, but not in total brain (Fig. S2C-F). Additionally, we found increased levels of MFN2 in AVs and decreased levels of MFN2 in brain lysates (Fig. S2G-J), suggesting that the HUWE1-MFN2 mitophagy pathway may also be preferentially utilized in the absence of PINK1.

BCL2L13 is another reported mitophagy receptor protein [29], but changes in levels of this protein it did not reach significance in our proteomic analysis (Fig. S2A, B). Nevertheless, we were curious to compare BCL2L13 levels in brain as well as enrichment within autophagosomes in PINK1^-/-^ mice relative to control. Immunoblotting detected higher levels of BCL2L13 in total brain, and higher levels in both the AV fractions, indicating BCL2L13 is enriched in autophagosome cargos isolated from PINK1^-/-^ mice (Fig. S2K-N).

Given the evidence that multiple alternative mitophagy pathways are upregulated in PINK1^-/-^ mice, we tested to see whether there was any detectable difference in the engulfment of damaged mitochondria in the brain in this model. We immunoblotted for FIS1 as a marker for mitochondria targeted to mitophagy [30] in our AV preparations, and found that FIS1 levels were maintained, and in fact increased, within PINK1^-/-^ derived AVs (Fig. 2I-L). Together, these changes suggest that there is upregulation of multiple compensatory pathways in the brain in order to clear damaged mitochondria in the absence of PINK1.

### PINK1^-/-^ neurons exhibit delayed autophagosome degradation

Given the upregulation of compensatory pathways, we asked whether damaged mitochondria were cleared with the same efficiency in the absence of PINK1. We immunoblotted for the mitochondrial marker HSP60 in Bafilomycin A-treated and Antimycin A-treated neuronal lysates (Fig. 3A). In PINK1^-/-^ neurons, HSP60 levels remained higher following mitochondrial damage (Fig. 3B), and addition of Bafilomycin A did not further increase the levels of HSP60 after mitochondrial damage as it did in the control neurons (Fig. 3C). Therefore, despite evidence for upregulation of compensatory pathways, PINK1^-/-^ neurons exhibited impaired mitochondrial flux following mitochondrial damage.

**Figure 3.**
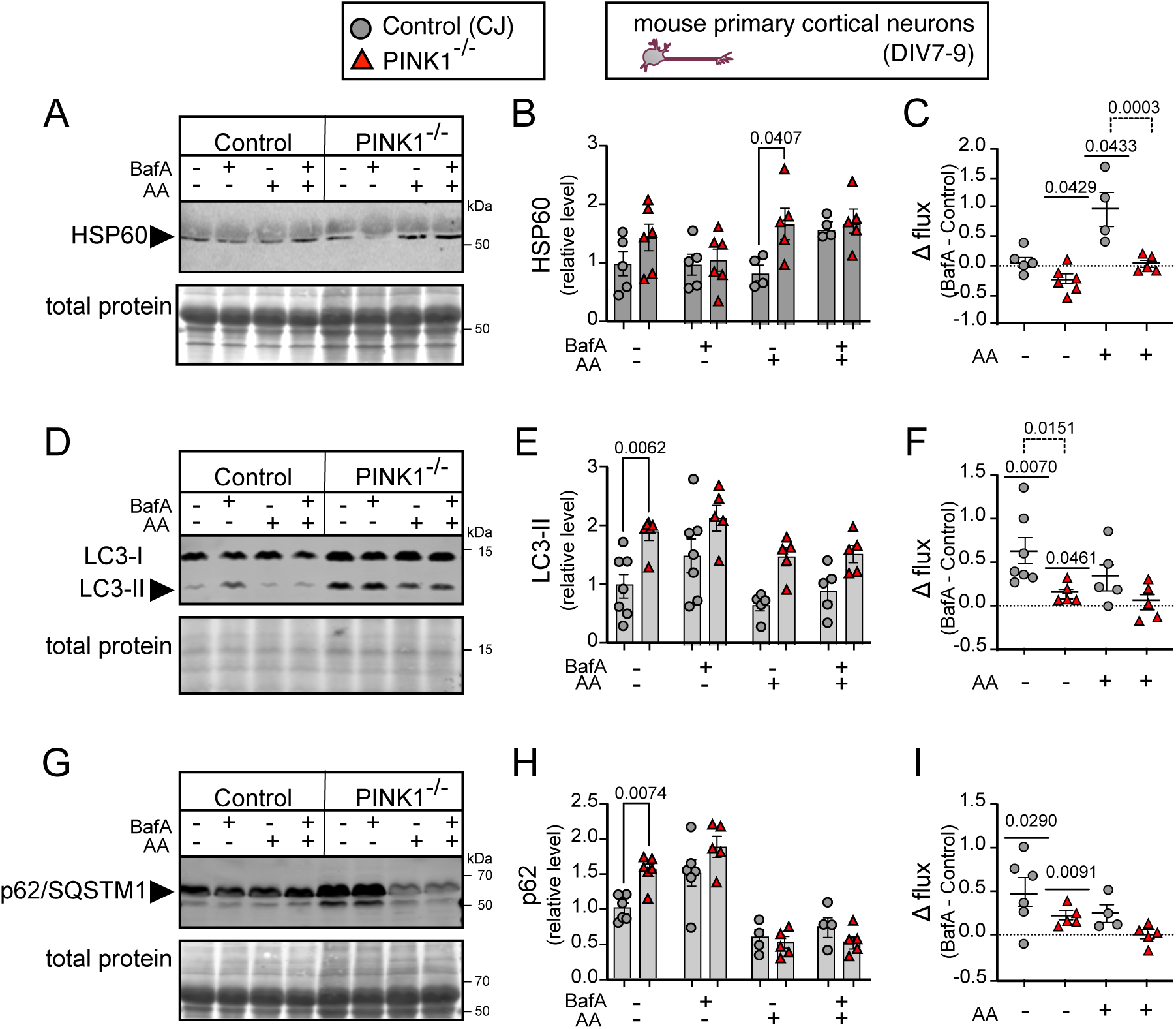
Autophagosome degradation is delayed in PINK1^-/-^ neurons **A-I.** Control and PINK1^-/-^ primary neurons were treated with Bafilomycin A (100nM for 2h) and Antimycin A (15 nM for 2 h) in a mitophagy flux assay, and cell lysates were probed by immunoblotting. **A, B.** Representative immunoblot (**A**) and quantification (**B**) (mean ± SEM, two way ANOVA with Šídák’s multiple comparisons test) of the levels of HSP60 in a mitophagy flux assay, normalized to total protein and control untreated levels. Control n ≥ 4, PINK1-/- n ≥ 5. **C.** Graph showing the change in flux levels of HSP60 from the paired Bafilomycin A treatment and untreated control, where 0 indicates no change in protein levels. One sample t and Wilcoxon test indicated by the line above the data points compares the mean ± SEM to the theoretical mean of 0. Dashed line between conditions indicates one way ANOVA with Šídák’s multiple comparisons test. Control n ≥ 4, PINK1^-/-^ n ≥ 5. **D, E.** Representative immunoblot (**D**) and quantifications (**E**) (mean ± SEM, two way ANOVA with Šídák’s multiple comparisons test) of LC3B-II levels in a mitophagy flux assay, normalized to total protein and control untreated levels. Control n ≥ 5, PINK1^-/-^ n ≥ 5. **F.** Graph showing the change in flux levels of LC3B-II from the paired Bafilomycin A treatment and untreated control, where 0 indicates no change in protein levels. One sample t and Wilcoxon test indicated by the line above the data points compares the mean ± SEM to the theoretical mean of 0. Dashed line between conditions indicates one way ANOVA with Šídák’s multiple comparisons test. Control n ≥ 5, PINK1^-/-^ n ≥ 5. **G, H.** Representative immunoblot (**G**) and quantifications (**H**) (mean ± SEM, two way ANOVA with Šídák’s multiple comparisons test) of p62/SQSTM1 levels in a mitophagy flux assay, normalized to total protein and control untreated levels. Control n ≥ 4, PINK1^-/-^ n ≥ 5. **I.** Graph showing the change in flux levels of p62/SQSTM1 from the paired Bafilomycin A treatment and untreated control, where 0 indicates no change in protein levels. One sample t and Wilcoxon test indicated by the line above the data points compares the mean ± SEM to the theoretical mean of 0. Control n ≥ 5, PINK1^-/-^ n ≥ 5.

Immunoblotting for LC3-II (Fig. 3D-F) and p62/SQSTM1 (Fig. 3G-I) in mitophagy flux assays suggests that the delay in mitochondrial degradation likely resulted from impaired autophagosome degradation, indicated by the higher levels of LC3-II and p62/SQSTM1 observed at baseline in PINK1^-/-^ mice (Fig. 3E, H); these increased levels in PINK1^-/-^ neurons do not further increase following Bafilomycin A treatment, suggesting there is a block in the degradative capacity of the lysosomes (Fig. 3F, I).

Together, proteomics, EM and immunoblotting results suggest that in a PINK1^-/-^ model of PD, alternative mitophagy pathways such as that mediated by the receptor BNIP3 are upregulated, although delayed degradation of autophagosomes was observed in flux assays. We hypothesize that this decreased flux may contribute to the accumulation of mitochondrial damage over time, leading to age-dependent neuronal loss.

### LRRK2^G2019S^-derived autophagosomes exhibit few changes in cargo selection, but show evidence of delayed maturation

While PINK1 mutations are rare and highly penetrant causes of PD, mutations in LRRK2, and in particular the LRRK2^G2019S^ mutation, are a common cause of hereditary PD [31]. Previously, we have shown that mutations in LRRK2 that induce hyperactive kinase activity lead to significant impairments in the trafficking and acidification of AVs, without affecting the formation or number of autophagosomes in primary neurons and human iPSC-derived neurons [25,32]. Thus, we were curious how autophagic cargos might be altered in the brains of LRRK2^G2019S^ mice, and how these alterations might compare to the changes observed in PINK1^-/-^ mice. We performed AV enrichment from genotype-matched control (B6NT) and LRRK2^G2019S^ mouse brain, in parallel to the studies on AVs isolated from the brain tissue of control and PINK1^-/-^ mice described previously (Fig. 4A). EM analysis of the isolated AV fractions indicated that >85% of vesicles were double membrane-bound autophagosomes, of which more than 50% contain mitochondria-like structures. In contrast to our findings from the PINK1^-/-^ model, we noted a ∼3- fold increase in the percent of more mature autolysosomes in LRRK2^G2019S^ -derived AVs (Fig. S3A,B), consistent with a delay in AV maturation in this model [25].

**Figure 4.**
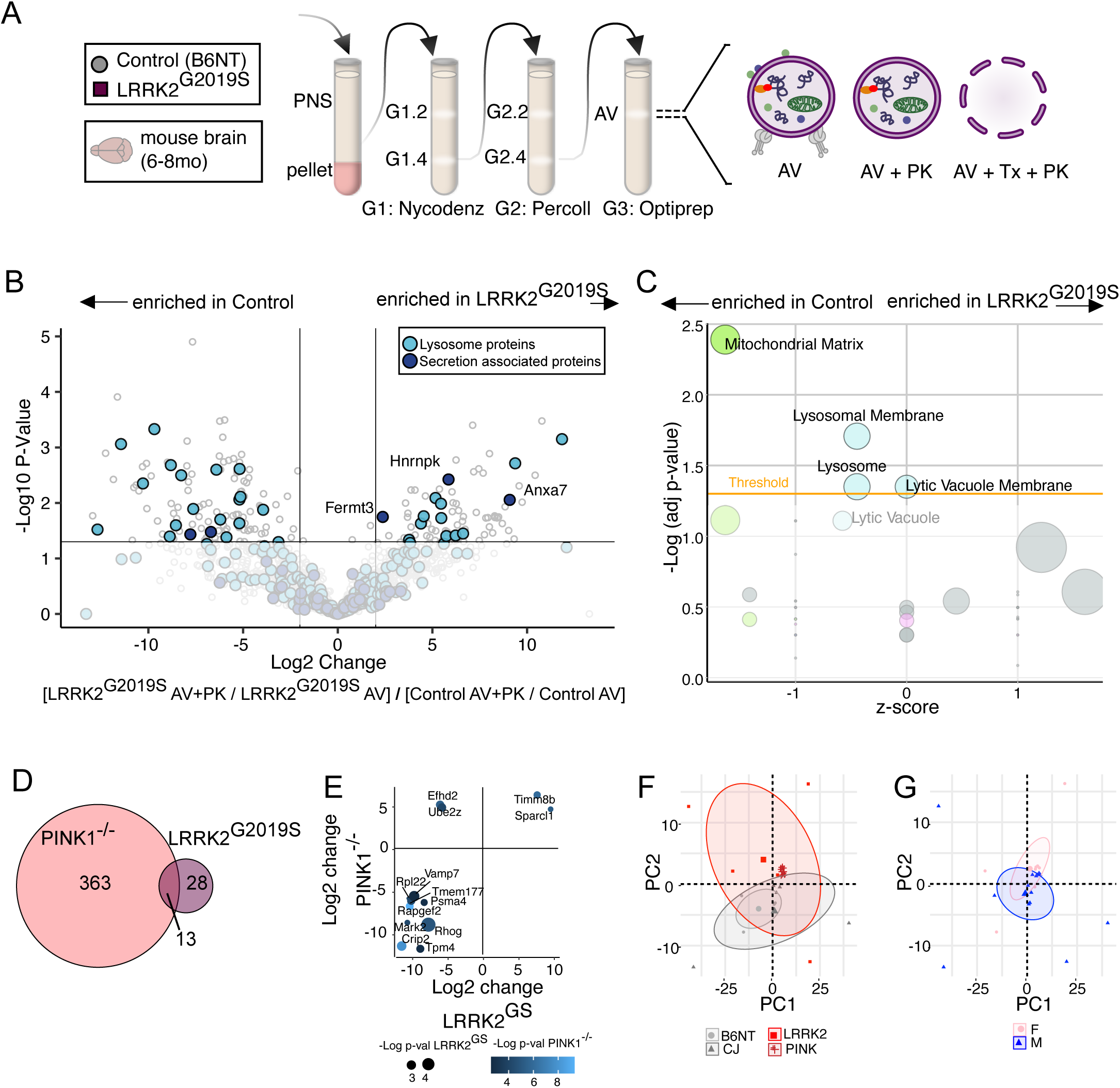
Characterization of LRRK^G2019S^ brain-derived autophagosome cargo indicates delays to autophagosome maturation **A.** Schematic representing the differential ultracentrifugation protocol and Proteinase-K digestion step for the enrichment of autophagic vesicles from control and LRRK2^G2019S^ mouse brain following euthanasia at 6-8 months of age. **B.** Volcano plot analysis of the ratio of AV+PK to AV fractions derived from the brain of LRRK2^G2019S^ relative to control mice. Black lines indicate significance thresholds of a p-value < 0.05 and a Log2 fold change > 2, and points are colored light blue if representing lysosomal proteins, and dark blue if representing secretion associated proteins. Control n = 4, LRRK2^G2019S^ n = 5. **C.** Bubble plot describing the significance and directionality of change of Gene Ontology terms from proteins whose cargo score was significantly changed between LRRK2 ^G2019S^ and control. Each bubble represents a unique GO term, colored light blue if representing a lysosomal term, magenta if representing a neuronal or synapse associated term, and green if representing a mitochondrial term. A significance threshold of an adjusted p-value <0.05 is marked in by a yellow line. The z-score represents the overall directionality of the cargo score, with a score of 0 indicating equal enrichment in both control and PINK1^-/-^. The size of the circle corresponds to the number of proteins identified in the GO term. **D.** Euler plot comparing the proteins with significantly changed cargo scores between PINK1^-/-^ and control and LRRK2^G2019S^ and control brain-derived AVs. **E.** Scatter plot showing the commonalities between significantly changed cargo scores from PINK1^-/-^ and control and LRRK2^G2019S^ and control brain-derived AVs. Size of circle denotes the LRRK2^G2019S^ p-value and color of circle denotes the PINK1^-/-^ p-value. **F, G.** Principal component analysis of the cargo score from controls (CJ and B6NT), PINK1^-/-^ and LRRK2^G2019S^ brain-derived AVs, colored by (**F**) genotype, or (**G**) sex.

Proteomic analysis of brain-derived AV and PK fractions from control and LRRK2^G2019S^ mice (B6NT n=4; LRRK2^G2019S^ n=5) highlighted fewer overall changes in raw abundances between the genotypes, with both increases and decreases in specific cargos observed in the LRRK2 model (Fig. S4A-D; Table S4, S5). Comparisons of cargo ratios and GO analysis indicate that proteins associated with the lysosome were decreased in LRRK2^G2019S^ relative to control mice (Fig. 3B, C; Table S6). Of note, this decrease in the cargo ratio is a result of increased lysosomal proteins in the AV fraction without the same magnitude increase in the PK fraction (Fig. S4C-F). As autophagosome-lysosome fusion results in lysosome membrane proteins on the external face of the autophagosome, these findings suggest autophagosome-lysosome fusion is maintained, but degradation is inefficient in the LRRK2 model, again consistent with the slowed maturation of AVs previously reported.

Next, we directly compared the changes in the AV and PK fractions from the PINK1^-/-^ and LRRK2^G2019S^ models of PD examined here. Approximately a third of the significant changes in the cargo ratio of LRRK2^G2019S^ AVs (13 out of 41) were common to the changes observed in the PINK1^-/-^ cargo ratio (Fig. 4D), with the directionality of change matched for 11 out of the 13 proteins (Fig. 4E). The majority of significantly changed peptides in the LRRK2^G2019S^ -derived AV and AV+PK fractions were also significantly changed in the PINK1^-/-^ AV and AV+PK fractions (Fig. S4G-J). Principal component analysis (PCA) on the cargo ratio of all four groups (CJ, B6NT, PINK1^-/-^ and LRRK2^G2019S^) indicated that controls and PD models tended to cluster separately along the second principal component axis (Fig. 4F), and sex did not contribute to either PC1 or PC2 (Fig. 4G). When PCA was performed on the peptide abundance values for the AV and PK fractions, the PINK1^-/-^ genotype clustered separately from the other groups along the first principal component axis, likely because of the increased peptide abundance. The second principal component axis clustered by sex (Fig. S4K-N).

### LRRK2^G2019S^ mice exhibit increased secretion of extracellular vesicles and decreased levels of PIKFYVE

Recent research has identified a causal relationship between lysosomal impairment and increased secretion of extracellular vesicles *in vitro* [33,34]. GO analysis of the significantly changed AV cargo proteins from LRRK2^G2019S^ brains found that terms for secretory or specific granules, both types of extracellularly secreted vesicles, were enriched (Fig. S4E, F). Specifically, we identified hnRNPK, associated with autophagy dependent secretion [35], FERMT3, a neuron-derived exosome marker associated with neurodegenerative disease [36], and ANXA7, which promotes calcium-dependent plasma membrane fusion and has been linked to regulating autophagy [37,38], all with increased cargo ratios in LRRK2^G2019S^ brain-derived AVs (Fig. 4B).

These observations led us to ask whether LRRK2^G2019S^ neurons exhibited increased secretion of extracellular vesicles (EVs), marked by TSG101 [39]. Levels of TSG101 were reduced in brain lysates from LRRK2^G2019S^ mice, while the low levels of TSG101 associated with AVs were unchanged from control mice (Figure 5A-D). We examined TSG101 secretion in primary cortical neurons from the LRRK2^G2019S^ model and found increased levels of TSG101 in conditioned media (Fig. 5E, F) without increases in the total cellular levels (Fig. S5A, B); importantly, elevated secretion of TSG101 could be reduced by inhibition of LRRK2 kinase activity with MLi-2. Human iPSC-derived glutamatergic LRRK2^G2019S^ knock-in neurons [25] also exhibited increased secretion of TSG101 into conditioned media, and again this increased secretion was reduced upon LRRK2 kinase inhibition with MLi-2 (Fig. 5G, Fig. S5C-E).

**Figure 5.**
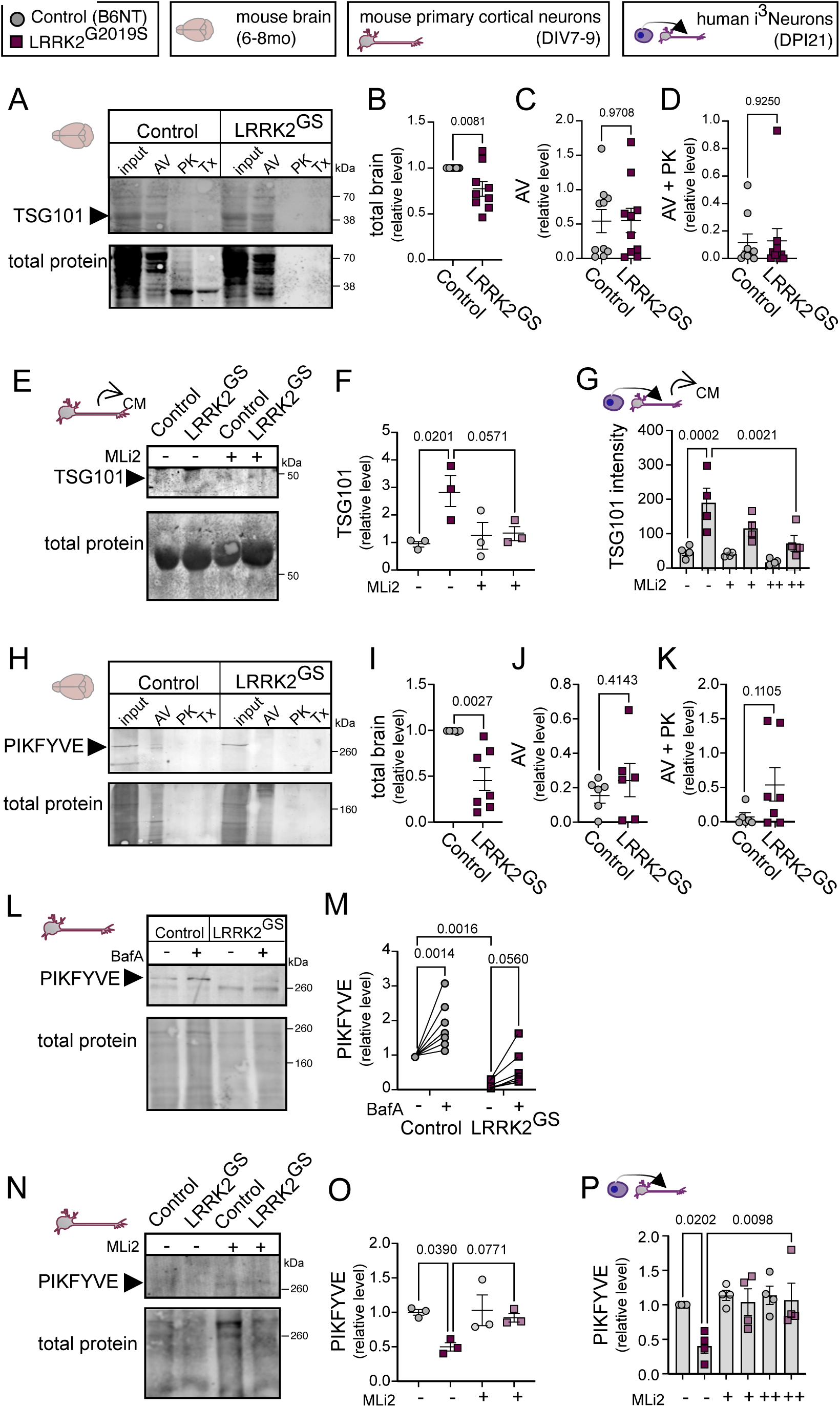
Increased extracellular vesicle secretion occurs in LRRK2^G2019S^ neurons **A-D.** Representative immunoblot (**A**) and quantifications (mean ± SEM, unpaired t test) of the levels of TSG101 in control and LRRK2^G2019S^ (**B**) total brain, (**C**) AV fraction, and (**D**) AV+PK fraction lysate, normalized to total protein and control total brain levels. Control n = 10, LRRK2^G2019S^ n = 10. **E, F.** Representative immunoblot (**E**) and quantification (**F**) (mean ± SEM, one way ANOVA with Šídák’s multiple comparisons test) of TSG101 levels in conditioned media from primary control and LRRK2^G2019S^ neurons, treated with DMSO control or MLi-2 (100 nM for 72 h), normalized to total protein and untreated control samples. Control n = 3, LRRK2^G2019S^ n = 3. **G.** Quantification (mean ± SEM, one way ANOVA with Šídák’s multiple comparisons test) of TSG101 levels in conditioned media from human iPSC-derived control and LRRK2^G2019S^ neurons, treated with DMSO control or MLi-2 (100 nM [+] or 1 μM [++] for 72 h), normalized to total protein. Control n = 4, LRRK2^G2019S^ n = 4. **H-K.** Representative immunoblot (**H**) and quantifications (mean ± SEM, unpaired t test) of the levels of PIKFYVE in control and LRRK2^G2019S^ (**I**) total brain, (**J**) AV fraction, and (**K**) AV+PK fraction lysate, normalized to total protein and control total brain levels. Control n = 6, LRRK2^G2019S^ n ≥ 6. **L, M.** Representative immunoblot (**L**) and quantifications (**M**) (mean ± SEM, two-way ANOVA with Fishers LSD) of PIKFYVE levels from primary control and LRRK2^G2019S^ neurons, treated with DMSO control or Bafilomycin A (100 nM for 2 h), normalized to total protein and untreated control samples. Control n = 7, LRRK2^G2019S^ n = 6. **N, O.** Representative immunoblot (**N**) and quantification (**O**) (mean ± SEM, one way ANOVA with with Šídák’s multiple comparisons test) of PIKFYVE levels in cell lysate from primary control and LRRK2^G2019S^ neurons, treated with DMSO control or MLi-2 (100 nM for 72 h), normalized to total protein and untreated control samples. Control n = 3, LRRK2^G2019S^ n = 3. **P.** Quantification (mean ± SEM, one way ANOVA with Šídák’s multiple comparisons test) of PIKFYVE levels in cell lysate from human iPSC-derived control and LRRK2^G2019S^ neurons, treated with DMSO control or MLi-2 (100 nM [+] or 1 μM [++] for 72 h), normalized to total protein and untreated control samples. Control n = 4, LRRK2^G2019S^ n = 4.

PIKFYVE is a kinase that phosphorylates PI3P to PI(3,5)P2, thereby regulating vesicular dynamics, lysosome fission and the fusion of multivesicular bodies to the lysosome [40–42]. Inhibition of PIKFYVE results in increased EV release [43,44]. We immunoblotted for PIKFYVE in total brain lysate, AV and PK fractions from LRRK2^G2019S^ and control mice (Fig. 5H). We found decreased levels of PIKFYVE in total brain in the LRRK2 model (Fig. 5I), while levels associated with the AV fractions remained unchanged (Fig. 5J, K). Based on these data, we wondered whether autophagy degrades PIKFYVE in neurons. We immunoblotted lysate from primary cortical neurons derived from control or LRRK2^G2019S^ mice treated with Bafilomycin A to prevent autophagosome degradation (Fig. 5L). Primary LRRK2^G2019S^ neurons also had reduced levels of PIKFYVE at baseline, and 2h of Bafilomycin A treatment increased the amount of PIKFYVE in both control and LRRK2^G2019S^ neurons (Fig. 5M).

We next investigated whether the reduced levels of PIKFYVE were a result of increased LRRK2 kinase activity. We found that MLi-2 rescued the levels of PIKFYVE in both primary cortical neurons (Fig. 5N, O) and iPSC-derived glutamatergic neurons with the LRRK2^G2019S^ mutation (Fig. 5P, Fig. S5F). Integrating our results with the previously published literature, we propose that lower levels of PIKFYVE as a result of increased LRRK2 kinase activity contribute to the increased secretion of EVs.

### LRRK2^G2019S^ increases the secretion of autophagy cargo TFAM and α-synuclein

hnRNPK, an RNA-binding protein known to be secreted in an autophagy dependent manner [45], had a higher cargo ratio in the LRRK2^G20129S^ AVs and thus we measured the levels in total brain, AVs and the secretion of hnRNPK from neurons in the LRRK2^G2019S^ model. We immunoblotted for hnRNPK in total brain lysate and AV fractions, but did not detect a significant change in hnRNPK levels in any of the fractions (Fig. S6A-D). However, we did observe increased secretion of hnRNPK into the conditioned media from the LRRK2^G2019S^ neurons that was mitigated by the addition of MLi-2 (Fig. 6A, B) while total intracellular protein levels remained unchanged (Fig. S6E, F). We saw a similar result analyzing conditioned media from LRRK2^G2019S^ human iPSC-derived neurons: we found increased secretion of hnRNPK from LRRK2^G2019S^ neurons as compared to isogenic control neurons, which was mitigated by treatment with MLi-2 (Fig. 6C, Fig. S6G-H).

**Figure 6.**
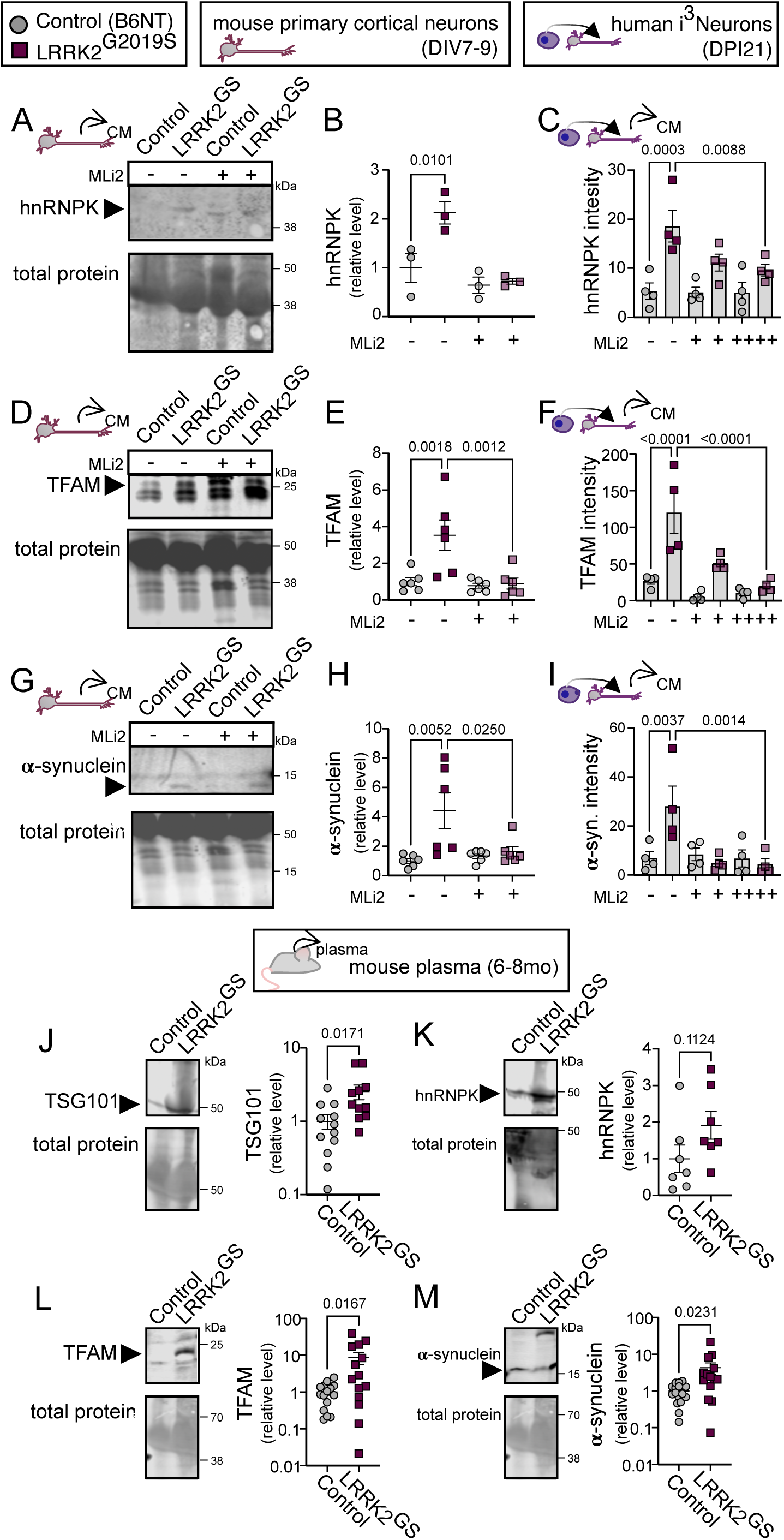
Increased autophagy-dependent secretion and secretion of autophagy cargos occurs in LRRK2^G2019S^ neurons **A, B.** Representative immunoblot (**A**) and quantification (**B**) (mean ± SEM, one way ANOVA with Šídák’s multiple comparisons test) of hnRNPK levels in conditioned media from primary control and LRRK2^G2019S^ neurons, treated with DMSO control or MLi-2 (100 nM for 72 h), normalized to total protein and untreated control samples. Control n = 3, LRRK2^G2019S^ n = 3. **C.** Quantification (mean ± SEM, one way ANOVA with Šídák’s multiple comparisons test) of hnRNPK levels in conditioned media from human iPSC-derived control and LRRK2^G2019S^ neurons, treated with DMSO control or MLi-2 (100 nM [+] or 1 μM [++] for 72 h), normalized to total protein. Control n = 4, LRRK2^G2019S^ n = 4. **D, E.** Representative immunoblot (**D**) and quantification (**E**) (mean ± SEM, one way ANOVA with Šídák’s multiple comparisons test) of TFAM levels in conditioned media from primary control and LRRK2^G2019S^ neurons, treated with DMSO control or MLi-2 (100 nM for 72 h), normalized to total protein and untreated control samples. Control n = 6, LRRK2^G2019S^ n = 6. **F.** Quantification (mean ± SEM, one way ANOVA with Šídák’s multiple comparisons test) of TFAM levels in conditioned media from human iPSC- derived control and LRRK2^G2019S^ neurons, treated with DMSO control or MLi-2 (100 nM [+] or 1 μM [++] for 72 h), normalized to total protein. Control n = 4, LRRK2^G2019S^ n = 4. **G, H.** Representative immunoblot (**G**) and quantification (**H**) (mean ± SEM, one way ANOVA with Šídák’s multiple comparisons test) of α-synuclein levels in conditioned media from primary control and LRRK2^G2019S^ neurons, treated with DMSO control or MLi-2 (100 nM for 72 h), normalized to total protein and untreated control samples. Control n = 6, LRRK2^G2019S^ n = 6. **I.** Quantification (mean ± SEM, one way ANOVA with Šídák’s multiple comparisons test) of α- synculein levels in conditioned media from human iPSC-derived control and LRRK2^G2019S^ neurons, treated with DMSO control or MLi-2 (100 nM [+] or 1 μM [++] for 72 h), normalized to total protein. Control n = 4, LRRK2^G2019S^ n = 4. **J-M.** Plasma from control and LRRK2^G2019S^ mice was collected and immunoblotted for (**J**) TSG101, (**K**) hnRNPK, (**L**) TFAM, and (**M**) α- synuclein. Quantification (mean ± SEM, unpaired t test), normalized to total protein and control samples, is shown. Control n ≥ 7, LRRK2^G2019S^ n ≥ 7.

Next, we looked for evidence of increased secretion of cargos associated with constitutive autophagy in the LRRK2^G2019S^ model. We immunoblotted for the presence of TFAM in brain-derived AVs and in conditioned media from neurons. While we saw no change in levels of TFAM in AV fractions from LRRK2^G2019S^ mice (Fig. S6J-M), release of TFAM into the conditioned media from primary cortical and iPSC-derived neurons was increased in a LRRK2 kinase activity-dependent manner (Fig. 6D-F, Fig. S6N-R).

α-synuclein is a synapse scaffolding protein and known autophagy cargo [19,46], while aggregated α-synuclein is a pathological hallmark of PD [47].The spread of aggregated or fibrillar α-synuclein in the brain correlates with disease progression, and it is thought to act in a prion-like manner that propagates neuronal dysfunction [48–54].Thus, we investigated whether α-synuclein levels were altered in LRRK2^G2019S^ derived AVs (Fig. S6S-V), and if there were increased levels of α-synuclein secretion in conditioned media from neurons in culture (Fig. 6G- I, Fig. S6W-AA). Levels of α-synuclein were reduced in AVs isolated from the brains of LRRK2^G2019S^ mice (Fig. S6U). However, no changes were detectable within the PK-protected fraction, relative to control (Fig. S6V). Consistent with our observations on hnRNPK and TFAM, secreted α-synuclein was increased from both primary cortical (Fig. 6G, H) and iPSC-derived neurons (Fig. 6I, Fig. S6Y) relative to control cells. Again, this increased secretion was a consequence of LRRK2 activity, as it was reversed by MLi-2 treatment of the neurons, and was not due to increases in intracellular protein levels (Fig. S6W, X, Z, AA).

Recently, there has been a clinical push for biomarker discovery for early detection of PD. To investigate whether the increased secretion of EVs and autophagy cargos could be detected in the circulating plasma of the LRRK2^G2019S^ mice, we compared the levels of TSG101, hnRNPK, TFAM and α-synuclein from the plasma of control and LRRK2^G2019S^ mice. We found increases in the plasma levels of TSG101, hnRNPK and autophagy cargo TFAM and α-synuclein in the LRKK2^G2019S^ plasma compared to control (Fig. 6J-M). Therefore, the increased secretion observed in neurons harboring hyperactive LRRK2 is detectable in circulating plasma.

### Inhibition of secretion in LRRK2^G2019S^ neurons results in cell death

To determine whether the increased secretion in the LRRK2^G2019S^ neurons was a compensatory change that was neuro-protective, we tested the sensitivity of control or LRRK2^G2019S^ primary cortical neurons to an inhibitor of the ESCRT-independent secretion pathway, GW4864. Overnight treatment with GW4864 effectively blocked secretion of EVs marked by TSG101 from both control and LRRK2^G2019S^ primary cortical neurons (Fig. 7A). LRRK2^G2019S^ neurons were significantly more sensitive to inhibition of secretion, as we observed increased levels of active caspase-3, a marker of cell death and apoptosis (Fig. 7B). In contrast, levels of NeuN, a neuronal transcription factor, were reduced (Fig. 7C). These observations suggest that the increased secretion of EVs observed in LRRK2^G2019S^ neurons is a compensatory mechanism that is critical to support neuronal health.

**Figure 7.**
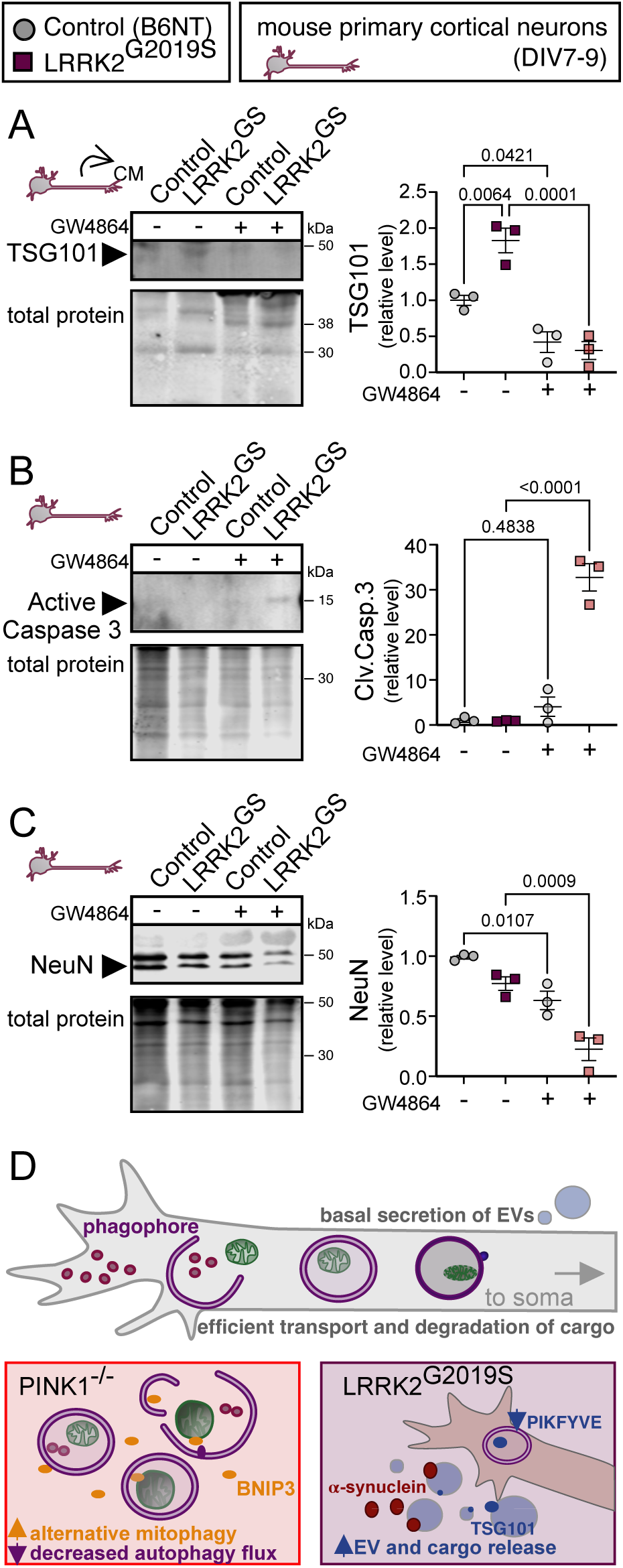
Inhibition of secretion in LRRK2^G2019S^ neurons results in cell death **A.** Representative immunoblot and quantification (mean ± SEM, one way ANOVA with Šídák’s multiple comparisons test) of TSG101 levels in conditioned media from primary control and LRRK2^G2019S^ neurons, treated with DMSO control or GW4864 (20 μM for 18 h), normalized to total protein and untreated control samples. Control n = 3, LRRK2^G2019S^ n = 3. **B.** Representative immunoblot and quantification (mean ± SEM, one way ANOVA with Šídák’s multiple comparisons test) of active caspase-3 levels in cell lysate from primary control and LRRK2^G2019S^ neurons, treated with DMSO control or GW4864 (20 μM for 18 h), normalized to total protein and untreated control samples. Control n = 3, LRRK2^G2019S^ n = 3. **C.** Representative immunoblot and quantification (mean ± SEM, one way ANOVA with Šídák’s multiple comparisons test) of NeuN levels in cell lysate from primary control and LRRK2^G2019S^ neurons, treated with DMSO control or GW4864 (20 μM for 18 h), normalized to total protein and untreated control samples. Control n = 3, LRRK2^G2019S^ n = 3. **D.** Cartoon schematic of key findings of increased alternative mitophagy in PINK1^-/-^ neurons and increased secretion in LRRK2^G2019S^ neurons.

## DISCUSSION

Proteomic analysis of autophagosomes has emerged as a powerful tool that provides insight into the role of autophagy to maintain neuronal health, and how this pathway is employed in development and disrupted in disease [11,55–57]. Here, we compared the proteomes of AVs isolated from two different models of PD and identified distinct autophagic signatures. We found upregulated pathways that complement the impairments introduced by the PD-associated mutations. Specifically, we observed increased levels of alternative mitophagy receptors engulfed within PINK1 null mice, and increased secretion of EVs and known autophagy cargo as a result of hyperactive LRRK2 activity in the LRRK2^G2019S^ model (Fig. 7D). We propose that these compensatory pathways maintain neuron health in the short term, and therefore may be responsible, at least in part, for the relatively mild phenotypes observed for either mouse model examined here. However, our results suggest that repercussions to the engagement of compensatory mechanisms may eventually emerge over longer timescales, and ultimately contribute to PD onset with aging. Specifically, these changes would be predicted to lead to delayed clearance of damaged mitochondria and/or increased secretion of α-synuclein or pro-inflammatory molecules such as mtDNA.

In the PINK1^-/-^ model, we found that mitochondria remained a major autophagy cargo under basal conditions, although we observed a lower proportion of AVs containing mitochondria. Strikingly, we noted increased levels of the selective mitophagy receptor protein BNIP3, which was degraded via mitophagy. However, the alternate mitophagy pathways engaged cannot compensate for the slow rate of degradation following mitochondrial damage in the absence of PINK1. We therefore predict that damaged mitochondria maybe engulfed rapidly but persist for longer within autophagosomes because of impaired autophagic flux in the absence of PINK1, and that detectable changes to mitochondrial health and neuronal health will only become apparent with accumulation over time, or upon severe mitochondrial stress. Indeed, this is consistent with the previous findings from PINK^-/-^ mouse models, which report increased mitochondrial dysfunction with age or upon induction of severe mitochondrial stress [58,59].

Fewer significant changes were detected in LRRK2^G2019S^ AVs compared to AVs isolated from PINK1^-/-^ mice, suggesting that cargo selection is not as dramatically altered in the LRRK2^G2019S^ model of PD. This observation is consistent with the known biology of LRRK2 as an important regulator of autophagosome trafficking and lysosome homeostasis, but not required for autophagosome cargo selection like PINK1. However, the changes to both lysosomal proteins and secretion-associated proteins that we observed in AVs from LRRK2^G2019S^ mice steered us to investigate changes to secretion. Analysis of conditioned media from cultured primary cortical neurons from LRRK2^G2019S^ mice, iPSC-induced human neurons gene-edited to express mutant LRRK2, as well as analysis of plasma from LRRK2 ^G2019S^ mice all demonstrate increased secretion of EVs, marked by increased levels of TSG101, and autophagy cargo including TFAM and α-synuclein. Further, the increased secretion observed from LRRK2-mutant neurons was reversed by inhibition of hyperactive kinase activity by MLi-2. Together, these data suggest that while cargo selection and uptake into autophagosomes are not profoundly affected in the LRRK2^G2019S^ model, there are changes consistent with an induction of secretory autophagy. This model fits with the recent findings that impaired lysosomal function and autophagosome-lysosome fusion increases the secretion of autophagosome cargo [33,43,60]. We propose that this increase in secretory autophagy is a compensatory mechanism to maintain proteostasis upon inhibition of lysosomal degradation, and we show that inhibition of secretion is detrimental to LRRK2^G2019S^ neuron survival.

The secretion of autophagy cargo observed here results in elevated circulating plasma levels of TSG101, hnRNPK, TFAM and α-synuclein. Therefore, this provides a tantalizing hypothesis that autophagy cargo may be useful to predict circulating biomarkers for the early detection of changes to lysosomal function or autophagic degradation, which are associated with early stages of neurodegeneration. Although they did not reach our p-value threshold, reported biomarkers for neurodegenerative disease Neurofilament Heavy Chain (NEFH) and CLSTN1 had cargo ratio Log2 changes of -5.97 and -11.71 respectively, while Neuregulin (NRGN), a predictive biomarker for Alzheimer disease progression, had a Log2 change of 13.28. Common autophagy cargo may be worthwhile targets for investigation.

We present a model in which decreased levels of PIKFYVE in the LRRK2^G2019S^ neurons promotes compensatory secretion of autophagy cargo to support neuron health. This pathway has been recently been reported to support the survival of neurons harboring an ALS mutation [61]. We hypothesize that the LRRK2 hyperactivity may somehow enhance PIKFYVE degradation, resulting in the lower baseline levels. We show that some of PIKFYVE turnover is dependent on autophagy, as levels accumulate following Bafilomycin A treatment. However, autophagic degradation of PIKFYVE to promote the compensatory secretion required for neuron health when autophagy is impaired introduces a serious vulnerability; if autophagy becomes further inhibited and there cannot be efficient compensation by the proteosome, PIKFYVE may accumulate and block secretion. Whether PIKFYVE will be a valuable targeted therapeutic for PD may depend on the status of LRRK2 activity or the levels of autophagic flux.

Although our data points to decreased PIKFYVE as a mechanism of increased secretion in the LRRK2^G2019S^ model, this may not be the only secretory pathway upregulated by hyperactive LRRK2. One illustrative example is the observed increase in α-synuclein secretion. If α- synuclein is secreted within or associated with EVs [62,63], the decrease in PIKFYVE levels may contribute to its release. Alternatively, increased secretion of α-synuclein has been reported in the context of the LRRK2^G2019S^ mutation via Rab35 [64,65]. A third potential mechanism is that the impaired acidification of autophagosomes contributes to the unconventional secretion of

α-synuclein, similar to the secretion of α-synuclein and autophagosome cargo via Rab27a mediated pathways [33,66]. While compensatory increases in secretion may maintain neuronal health in the short term, we propose that this increased secretion of cargos such as α-synuclein or TFAM may ultimately prove detrimental in the context of PD progression. We found increased secretion of monomeric 14 kDa α-synuclein, but also a seemingly covalent dimeric form at 28 kDa, similar to the nonfibrillar oligomer previously described [67]. As different oligomers of α-synuclein have different effects on cell signaling and Lewy body formation, secretion of multiple molecular weight oligomers of α-synuclein may have pleiotropic effects on neighboring neurons [68]. If LRRK2 mutations cause monomeric or nonfibrillar oligomeric secretion of α-synuclein secretion, it may prevent intra-neuronal buildup preceding pathogenic aggregation [69]. Indeed, decreased α-synuclein in the CSF is correlated with PD-diagnosis, suggesting that secretion may be protective [70]. However if the α-synuclein is aggregated prior to secretion, as is the case with pre-formed fibrils, secretion in response to impaired autophagy could propagate PD pathology and accelerate disease progression as previously found [71,72]. Similarly, if TFAM and potentially mitochondrial DNA is being secreted, might this promote an inflammatory response that contributes to the worsening of PD progression?

In all, our proteomic analysis identifies changes to the autophagy pathway associated with PD, namely pathways that compensate as a response to the germline mutation. We predict that the adaptive changes delay, but do not prevent, neurodegenerative disease progression, and may ultimately contribute to mitochondrial dysfunction, neuroinflammation, or Lewy body propagation. Our findings further cement the crucial role of autophagy in maintaining neuron health and demonstrate the importance of this pathway in neurodegenerative disease progression.

## Acknowledgments

We thank Karen Wallace Jahn for assistance with animal models, Sierra Palumbos and Bishal Basak for assistance in primary neuron dissections, and J.A. Paulo for proteomics support. We thank the Electron Microscopy Resource Lab for negative stain preparation for transmission electron microscopy.

This research was supported by the German Research Foundation (DFG; BO 5434/2-1 to C.A.B.), by National Institutes of Health grants NS083524 (JWH) and NS060698 (ELFH), and by Sloan Kettering Institute startup funds (AO). This research was funded in part by Aligning Science Across Parkinon’s ASAP-000350 (ELFH) and ASAP-000282 (JWH) through the Michael J. Fox Foundation for Parkinson’s Research (MJFF) and in part supported by the Memorial Sloan Kettering Cancer Center Support Grant P30CA008748 (A.O.). For the purpose of open access, the author has applied a CC BY public copyright license to all Author Accepted Manuscripts arising from this submission.

## Author Contributions

JG, ELFH and JWH, project design and conceptualization; JG and ELFH, co-writing – original draft; JG, ELFH, JWH, CAB, and AO, editing manuscript; CAB, assisted with primary cortical neuron isolation, culture and sample collection; MA, IB assistance; AO, proteomics data collection and analysis; JG, resources, data collection, data analysis and interpretation; ELFH, JWH funding acquisition.

## Inclusion and Diversity

We worked to ensure sex balance in the selection of rodent subjects. While citing references scientifically relevant for this work, we also actively worked to promote gender balance in our reference list.

## STAR Methods

### RESOURCE AVAILABILITY

#### Lead contact

Further information and requests for resources and reagents should be directed to and will be fulfilled by the lead contact, Erika Holzbaur.

#### Data and code availability

The MS proteomics data have been deposited to the MassIVE repository with the dataset identifier MSV000090264.

### EXPERIMENTAL MODEL AND SUBJECT DETAILS

#### Animal models

For mass spectrometry and immunoblotting analysis of brain-derived autophagic vesicles, the genotypes C57BL/6NTac (B6NT) control and C57BL/6-*Lrrk2^tm4.1Arte^* (LRRK2^G2019S^ knock-in) mice, available from Taconic Models #B6 and #13940 respectively, and C57BL/6J (CJ) control and B6.129S4-PINK1^tm1Shn/J^ (PINK1^-/-^) mice, available from Jackson Laboratories models #:000664 and #017946 respectively, were used. Mice of both sexes at 6-8 months of age were euthanized according to University of Pennsylvania Institutional Animal Care and Use Committee approved procedures and the brain above the brainstem was removed and homogenized in a sucrose buffer (see method details).

#### Primary cell cultures

Primary mouse cortical neurons were dissected and cultured in our laboratory. Briefly, E15.5 embryos from mouse lines described above were dissected and the cortex was removed and dissociated with 0.25% trypsin and trituration. Neurons were plated in attachment media (MEM supplemented with 10% horse serum, 33 mM D-glucose and 1 mM sodium pyruvate) on poly-L-lysine coated 35 mm dishes. After 4-6h, media was replaced with maintenance media (Neurobasal [GIBCO] supplemented with 2% B-27 [GIBCO], 33 mM D-glucose [Sigma], 2 mM GlutaMAX [GIBCO], 100 U/mL penicillin and 100 mg/mL streptomycin [Sigma]). AraC (1 μM) was added the day after plating to prevent glia cell proliferation. Every 3-4 days, 40% of the media was replaced with fresh Maintenance Media and at day in vitro (DIV) 7-9 the neurons were used for biochemical analysis. For MLi-2 treatment, neurons were incubated in 100 nM MLi-2 over 72 hours or DMSO control. Bafilomycin A treatment (100 nM) and Antimycin A treatment (15 nM) was 2 h. GW4864 treatment (20 μM) was 18 h.

#### Human iPSC-derived cell cultures

Description of the generation, culture, and differentiation of the iPSC line WTC11 control and LRRK2^G2019S^ knock-in has been previously described [25]. Briefly, human i3N iPSCs that harbor a doxycycline-inducible mNGN2 transgene in the AAVS1 safeharbor locus, a gift from M. Ward (National Institutes of Health, Maryland) were CRISPR-edited to knock-in the LRRK2^G2019S^ mutation. Cytogenetic analysis of G-banded metaphase cells demonstrated a normal male karyotype (Cell Line Genetics), and mycoplasma testing was negative. i3N iPSCs were cultured on Growth Factor Reduced Matrigel (Corning) coated plates and fed daily with mTeSR medium (StemCell). Differentiation into i3Neurons was performed following an established protocol [73]. i3N iPSCs were split with Accutase (Sigma) and plated on Matrigel-coated dishes in Induction Medium (DMEM/F12 containing 2 μg/mL doxycycline, 1% N2-supplement [Gibco], 1% NEAA [Gibco] and 1% GlutaMAX [Gibco]). After 3 days, pre-differentiated i3Neurons were dissociated with Accutase and cryo-preserved. On day of use, pre-differentiated i3Neurons were thawed and plated on poly-L-ornithine coated dishes at an appropriate density, ∼300,000 cells per 35mm dish. i3Neurons were cultured in BrainPhys Neuronal Medium (StemCell) supplemented with 2% B27 (Gibco), 10 ng/mL BDNF (PeproTech), 10 ng/mL NT-3 (PeproTech) and 1 μg/mL Laminin (Corning). Every 3–4 days, 40% of the medium was replaced with fresh culture medium. Biochemistry experiments were performed 18 days after thawing pre-differentiated i3Neurons (days post induction DPI21). For MLi-2 treatment, neurons were incubated in 100 nM or 1 μM MLi-2 over 72 hours or DMSO control.

### METHOD DETAILS

#### Isolation of autophagic vesicles by differential centrifugation

Enriched autophagosome fractions were isolated following a protocol modified from Strømhaug et al., 1998 and Maday et al., 2014. Briefly, one mouse brain was collected in a 250mM sucrose solution buffered with 10 μM HEPES and 1mM EDTA at pH 7.3, homogenized using a tissue grinder, incubated with Gly-Phe-β-naphthylamide (GPN) for 7 min at 37°C to destroy lysosomes and subsequently subjected to three differential centrifugations through 9.5% Nycodenz and 33% Percoll and 30% Optiprep discontinuous gradients to isolate vesicles of the appropriate size and density. Following collection, the autophagic vesicle enriched fraction (AV) was divided into three, one third was treated with 10 μg Proteinase K for 45min at 37°C, similar to Le Guerroué et al., 2017 and Zellner et al., 2021, to degrade non-membrane protected proteins and enrich for internal autophagosome cargo (AV+PK), one third was membrane permeabilized by the addition of 0.2% Triton X-100 prior to the same proteinase K treatment as a negative control (AP+Tx+PK), and the other third was left untreated for identification of all internal and externally-associated proteins on autophagosomes. AV-enriched fractions were subsequently used for mass spectrometry, electron microscopy, and immunoblotting.

#### Proteomics – sample preparation and digestion

The AV and AV+PK fractions from independent mouse brain preparations were lysed with RIPA buffer (50 mM HEPES (pH 7.4), 150 mM NaCl, 1% sodium deoxycholate, 1% NP-40, 0.1% SDS, 2.5 mM MgCl2, 10 mM sodium glycerophosphate, 10 mM sodium biphosphate) containing 1 µg/ml aprotinin, 1 µg/ml leupeptin, 1 mM benzamidine, 1 mM AEBSF and 1% final SDS. Lysates were sonicated on ice three times, followed by centrifugation (13000 rpm, 5 min). Protein concentration was measured by Bradford assay. Protein extracts (50 ug) were subjected to disulfide bond reduction with 5 mM TCEP (room temperature, 10 min) and alkylation with 25 mM chloroacetamide (room temperature, 20 min) and followed by TCA precipitation, prior to protease digestion. Samples were resuspended in 100 mM EPPS, pH 8.5 containing 0.1% RapiGest and digested at 37°C for 8 h with Trypsin at a 100:1 protein-to-protease ratio. Trypsin was then added at a 100:1 protein-to-protease ratio and the reaction was incubated for 6 h at 37°C. Following incubation, digestion efficiency of a small aliquot was tested. The sample was vacuum centrifuged to near dryness, resuspended in 5% formic acid for 15 min, centrifuged at 10000×g for 5 minutes at room temperature and subjected to subjected to C18 StageTip desalting.

#### Proteomics – Liquid chromatography and tandem mass spectrometry

Mass spectrometry data were collected using an Orbitrap Eclipse Tribrid Mass Spectrometer (Thermo Fisher Scientific), combined with a high-field asymmetric waveform ion mobility spectrometry (FAIMS) Pro interface, coupled to a Proxeon EASY-nLC1000 liquid chromatography (LC) pump (Thermo Fisher Scientific). Peptides were separated on a 100 μm inner diameter microcapillary column packed in house with ∼35 cm of Accucore150 resin (2.6 μm, 150 Å, Thermo Fisher Scientific, San Jose, CA) with a gradient (ACN, 0.1% FA) over a total 60 min run at ∼550 nL/min. For analysis, we loaded 1/4 of each fraction onto the column. The scan sequence began with an MS^1^ spectrum (Orbitrap analysis resolution 120,000 at 200 Th; mass range 375−1500 m/z; automatic gain control (AGC) target 4×10^5^; maximum injection time 50 ms) and peak-picking algorithm Advanced Peak Determination was used. Precursors for MS^2^ analysis were selected using a cycle type of 1 sec/CV method (FAIMS CV=-40/-60/-80 [74]). MS^2^ analysis consisted of collision-induced dissociation (quadrupole ion trap analysis; Rapid scan rate; AGC 2.0×10^4^; isolation window 0.7 Th; normalized collision energy (NCE) 35; maximum injection time 35 ms). Monoisotopic peak assignment was used, determined charge states between 2 and 6 were required for sequencing, previously interrogated precursors were excluded using a dynamic window (60 s ± 10 ppm) and dependent scan was performed on a single charge state per precursor.

#### Proteomics - Data analysis

Mass spectra were processed using Protein Discoverer using the Minora algorithm (set to default parameters). Database searching included all canonical entries from the mouse Reference Proteome UniProt database (SwissProt – 2019-12), as well as an in-house curated list of contaminants. The identification of proteins was performed using the SEQUEST-HT engine against the database using the following parameters: a tolerance level of 10 ppm for MS^1^ and 0.6 Da for MS^2^ post-recalibration and the false discovery rate of the Percolator decoy database search was set to 1%. Trypsin was used as the digestion enzyme, two missed cleavages were allowed, and the minimal peptide length was set to 7 amino acids. Carbamidomethylation of cysteine residues (+57.021 Da) were set as static modifications, while oxidation of methionine residues (+15.995 Da) was set as a variable modification. Final protein-level FDR was set to 1%. Precursor abundance quantification was determined based on intensity and the minimum replicate feature parameter was set at 50%. Proteins were quantified based on unique and razor peptides.

Protein quantification values were exported for further analysis in Microsoft Excel and Perseus [75] and statistical test and parameters used are indicated in the corresponding Supplementary Data Tables datasets. Briefly, Welch’s t-test analysis was performed to compare two datasets, using s0 parameter (in essence a minimal fold change cut-off) and correction for multiple comparison was achieved by the permutation-based FDR method, both functions that are built-in in Perseus software.

#### Plasma collection

Blood from mice was collected following IACUC-approved euthanasia and decapitation. Approximately 200 μl of blood was collected in a microvette tube coated in EDTA, and the samples were spun at 2000 x g for 5 minutes at room temperature. 80μl of plasma was collected, diluted in 200 μl PBS and denaturing buffer was added to a 1x final concentration, then boiled for 5 min at 95°C.

#### Immunoblotting

Samples were lysed in RIPA buffer (50 mM Tris-HCl, 150 mM NaCl, 0.1% Triton X-100, 0.5% deoxycholate, 0.1% SDS, 2x Halt Protease and Phosphatase inhibitor, PMSF, Pepstatin A, TAME and Leupeptin), centrifuged at 18,000 x g for 20 min to clear unlysed and membranous fractions, and then the protein concentration was determined by Bradford assay. Conditioned media from DIV14 primary cortical neurons was first pulse spun to remove cell debris in the media (5000 x g for 2 min) before denaturing buffer was added, and boiled for 5 min at 95°C. Proteins were resolved on 6%, 8%, 10%, 12% or 15% SDS-PAGE gels, based on size of proteins to be identified. Proteins were transferred to Immobilon-FL PVDF membranes (Millipore) using a wet blot transfer system (BioRad). 15% gels were transferred in buffer containing 20% methanol. Membranes were stained for total protein using Li-Cor Revert Total Protein Stain. Following imaging, the total protein was destained, blocked for 5min at RT with EveryBlot blocking buffer (BioRad) and incubated with primary antibodies diluted in TrueBlack WB antibody diluent + 0.2% Tween-20 overnight at 4°C. Membranes were washed three times in TBS + 0.1% Tween-20 and incubated with secondary antibodies (1:20,000 dilution) in TrueBlack WB antibody diluent + 0.2% Tween-20 + 0.1% SDS for 1h at RT. Following three washes in TBS+ 0.1% Tween-20, membranes were imaged using Odyssey CLx Infrared Imaging System (LI-COR), and quantification of protein levels was performed using ImageStudio (Li-Cor).

For quantification of immunoblot data, an independent biological replicate is defined as a separate brain, cortical neuron preparation, or i3N differentiation. Data was excluded if the total protein levels were unquantifiable.

#### Electron microscopy

AV were pelleted and fixed with 2.5% glutaraldehyde, 2.0% paraformaldehyde in 0.1M sodium cacodylate buffer, pH 7.4, overnight at 4°C. Fixed samples were then transferred to the Electron Microscopy Resource Laboratory at the University of Pennsylvania, where all subsequent steps were performed. After buffer washes, the samples were post-fixed in 2.0% osmium tetroxide for 1 h at room temperature and then washed again in buffer, followed by dH_2_O. After dehydration through a graded ethanol series, the tissue was infiltrated and embedded in EMbed-812 (Electron Microscopy Sciences, Fort Washington, PA). Thin sections were stained with lead citrate and examined with a JEOL 1010 electron microscope fitted with a Hamamatsu digital camera and AMT Advantage image capture software. Regions of dense AVs were chosen for imaging. Biological replicates are defined as separate brain-derived AV preparations.

### QUANTIFICATION AND STATISTICAL ANALYSIS

Mitochondrial annotations and sublocalizations were performed using the MitoCarta3.0 and UniProt databases [76].

GraphPad prism software (v9.1.0) was used for statistical analysis. Generally, the statistical test performed for all immunoblot analyses is unpaired t test for comparisons of two categories, and Ordinary one-way or two-way ANOVA with Šídák’s multiple comparisons test for comparisons of three or more groups. The specific statistical test employed is listed in the figure legend. Biological replicates (n), defined in the method details, are always displayed as individual data points and the precision measures (mean ± SEM) are displayed. Significance was defined as a p-value < 0.05, and directly reported in the figure. R (v4.0.4) was used to generate volcano plots (GGplot package), Euler diagrams (VennDiagram package), XY plots of Log2 change of mass spectrometry data sets (GGplot package), GO bubble plots (GOplot package) and PCA analysis (FactoMineR package). Enrichr [77–79] and SynGO [80] were used to compute the p-values of gene ontology term enrichment. Cargo scores identified as significantly changed between groups by analysis on Perseus were input into the softwares – the p-value calculation is dependent on Fisher’s exact test and the q-value displays the Benjamini-Hochburg multiple hypothesis testing correction. Abundance ratios for the AV and AV+PK fractions were defined as significant if p-value was < 0.05 and the Log2 fold change was > 2. Enrichr precomputes a background expected rank for each term in the gene set library. Neither Enrichr nor SynGO software takes into account the background protein expression levels in specific organs.

### KEY RESOURCES TABLE

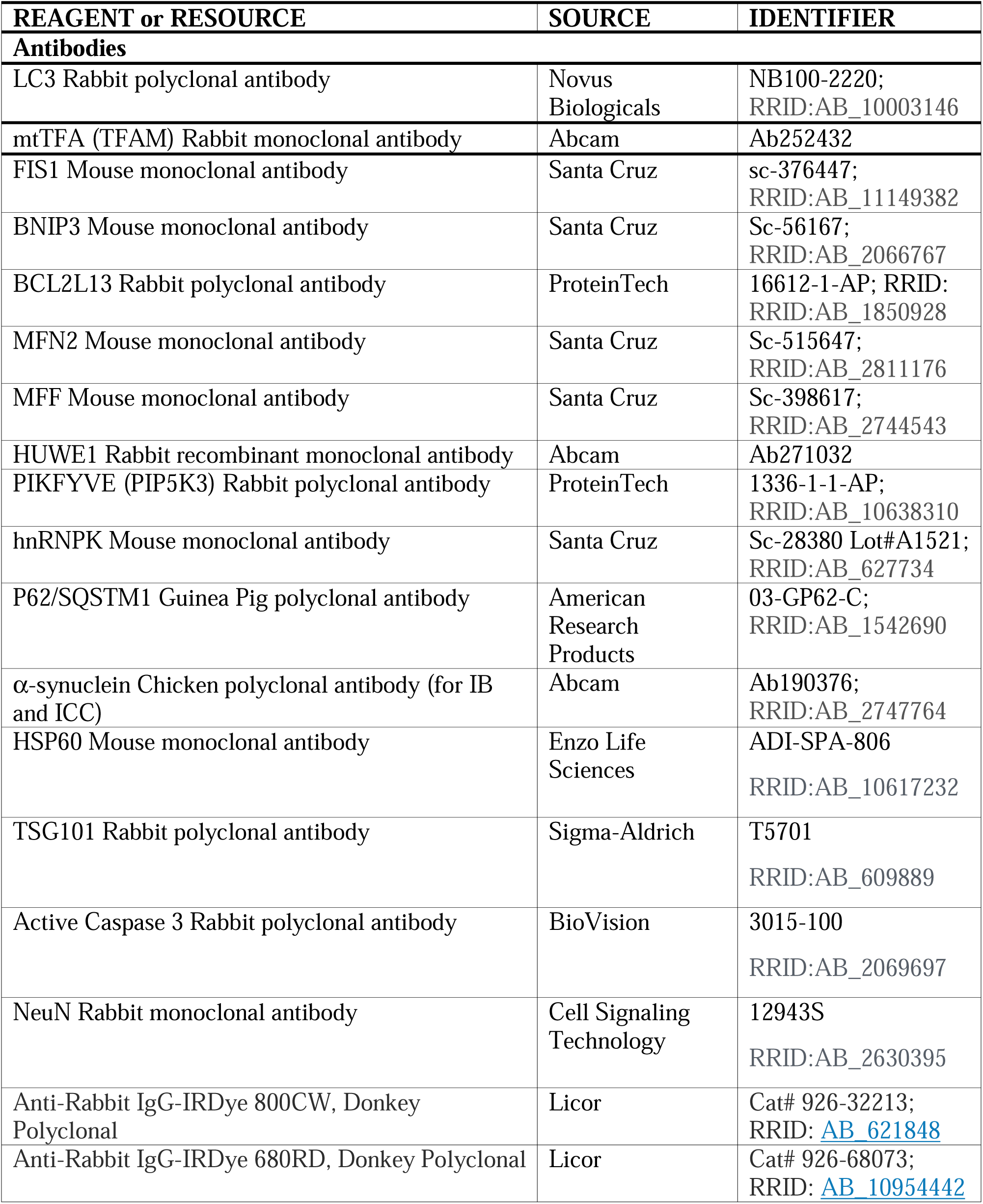
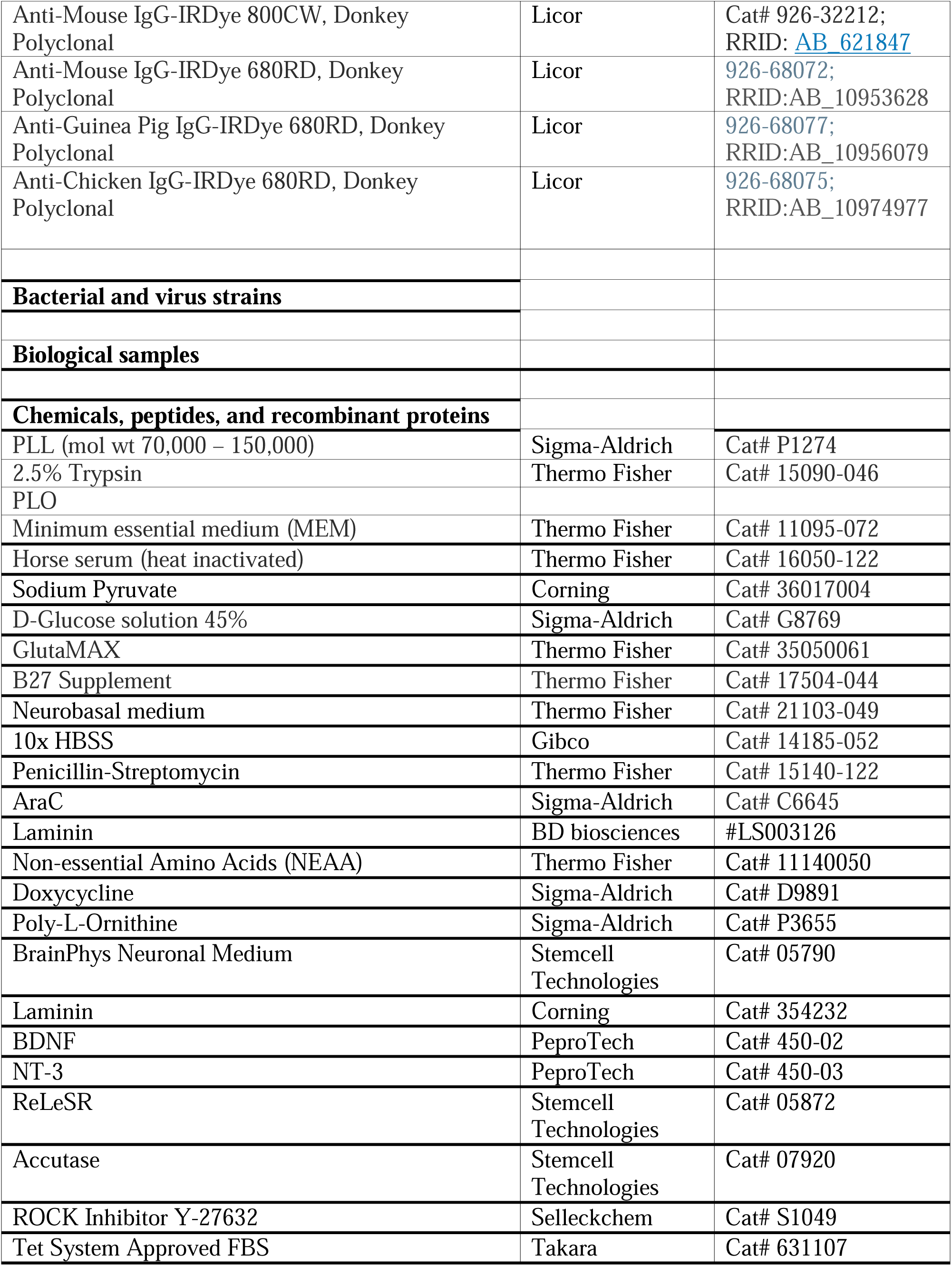
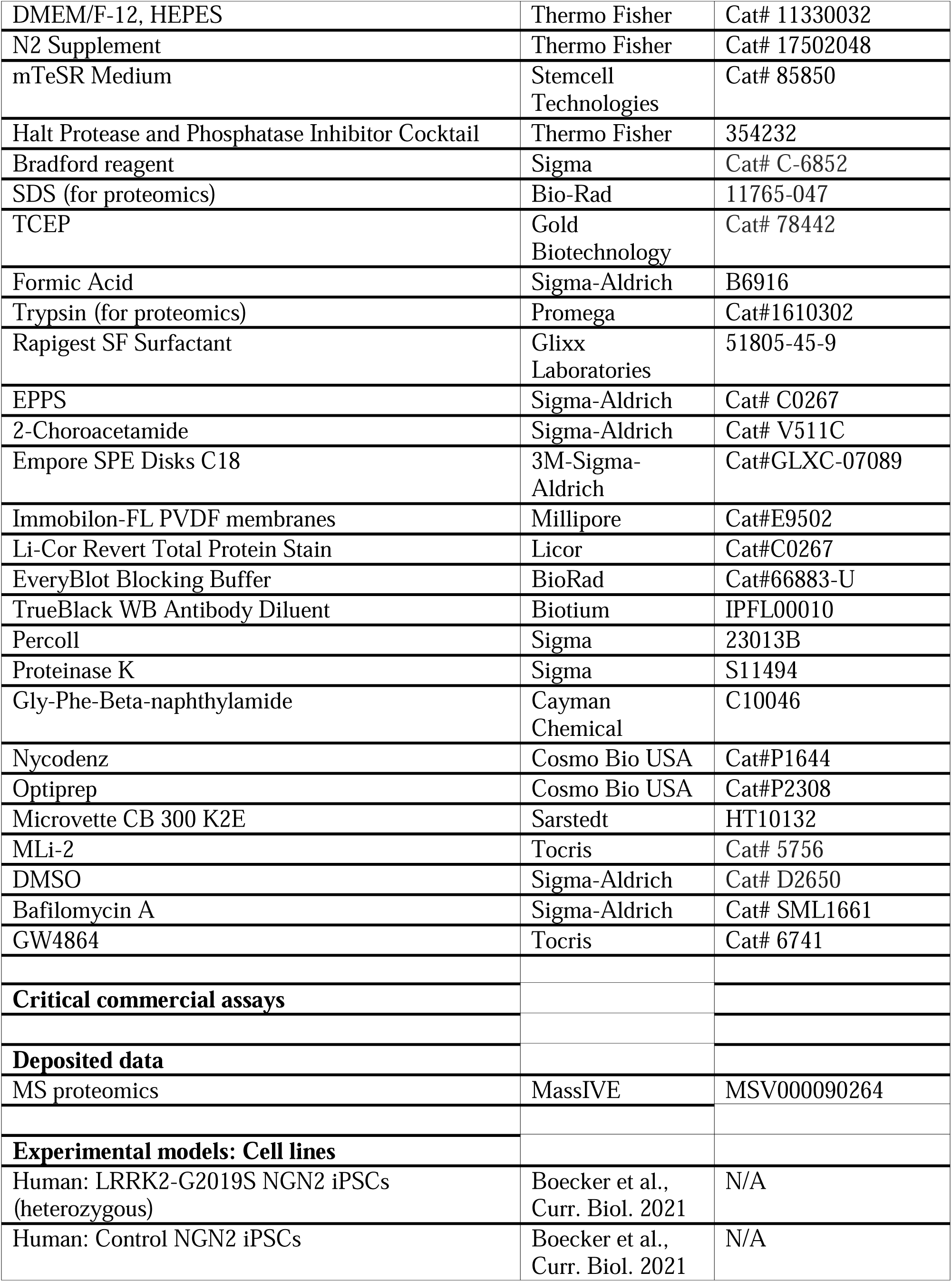
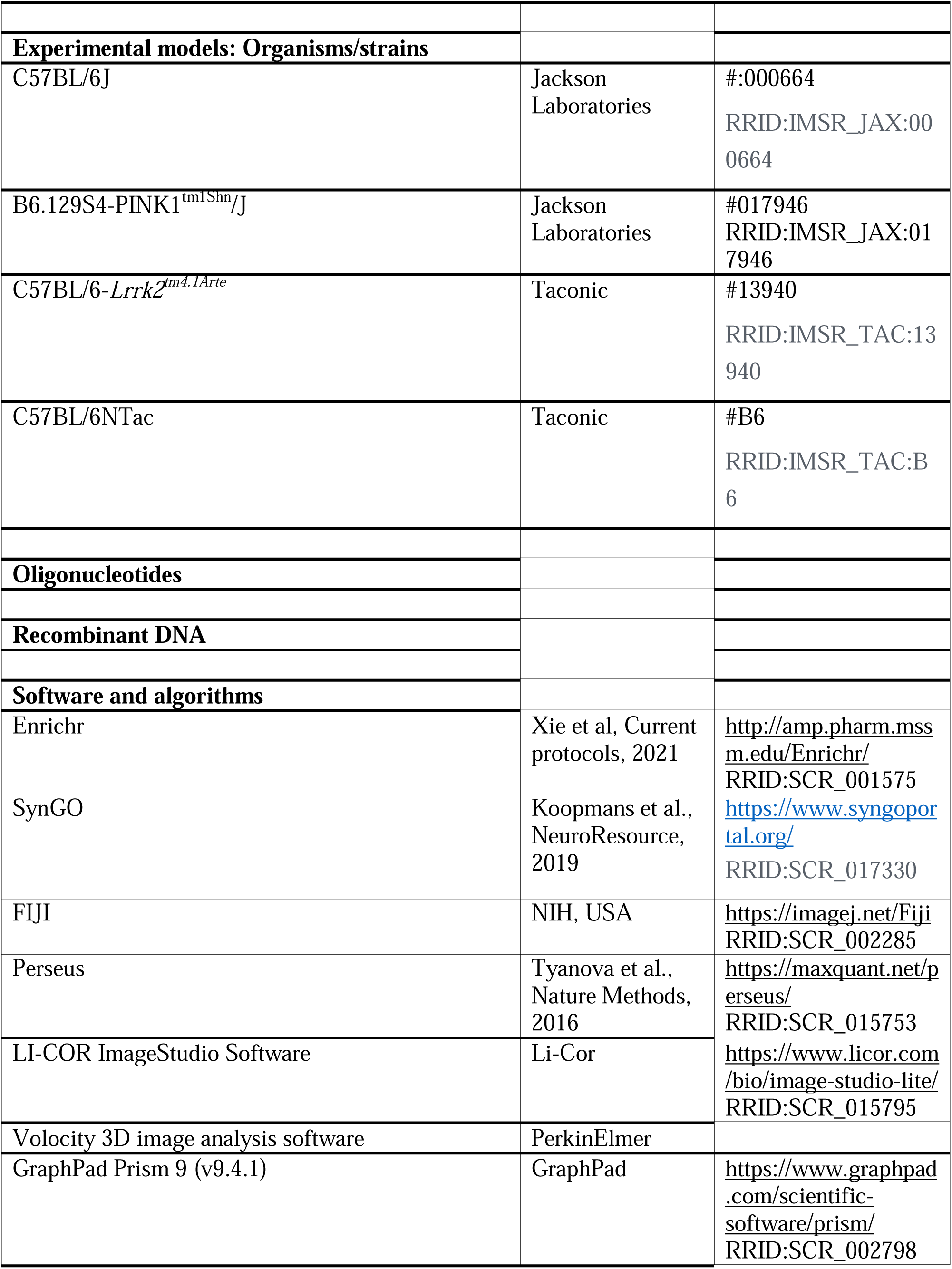
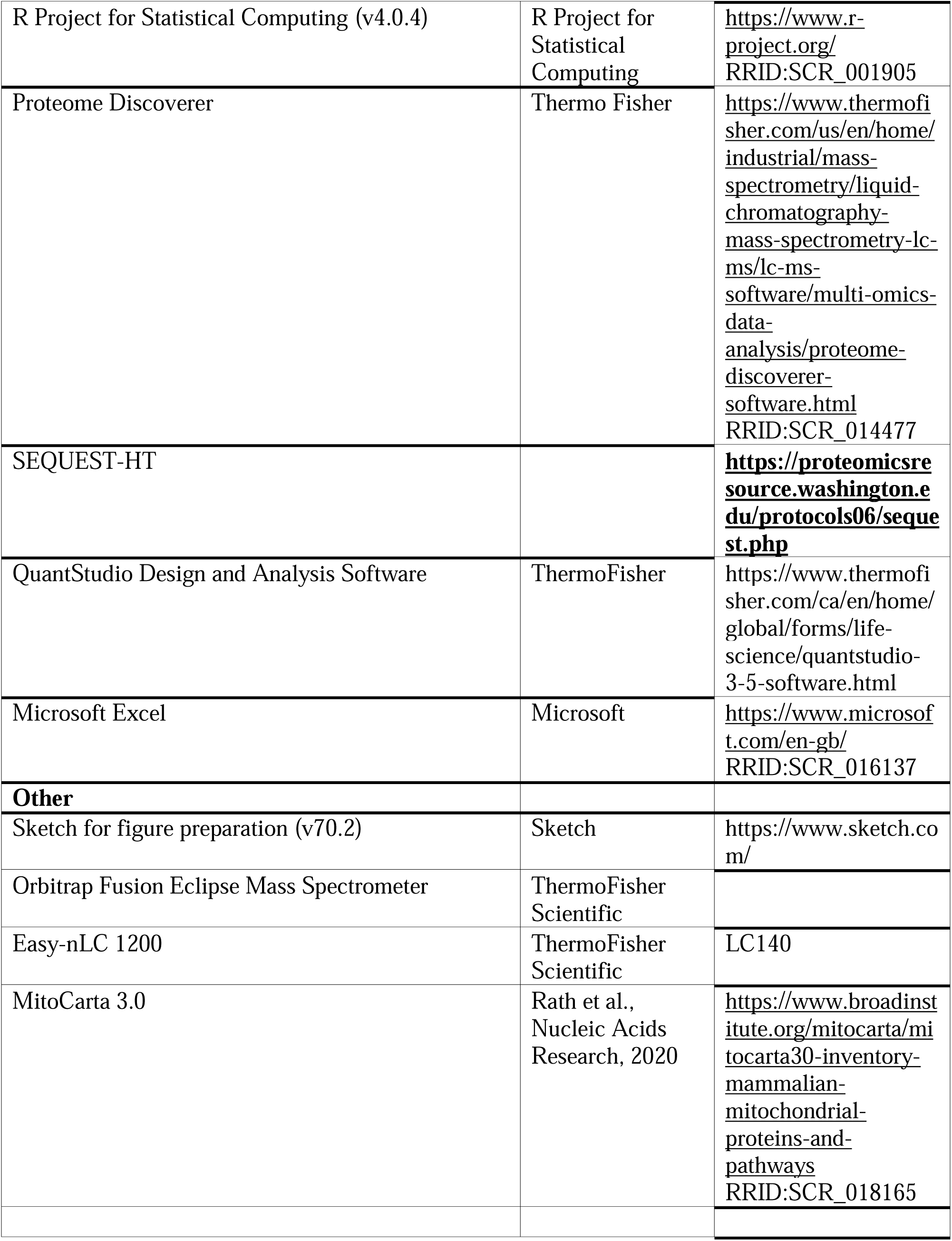

---

**Supplementary Figure 1.**
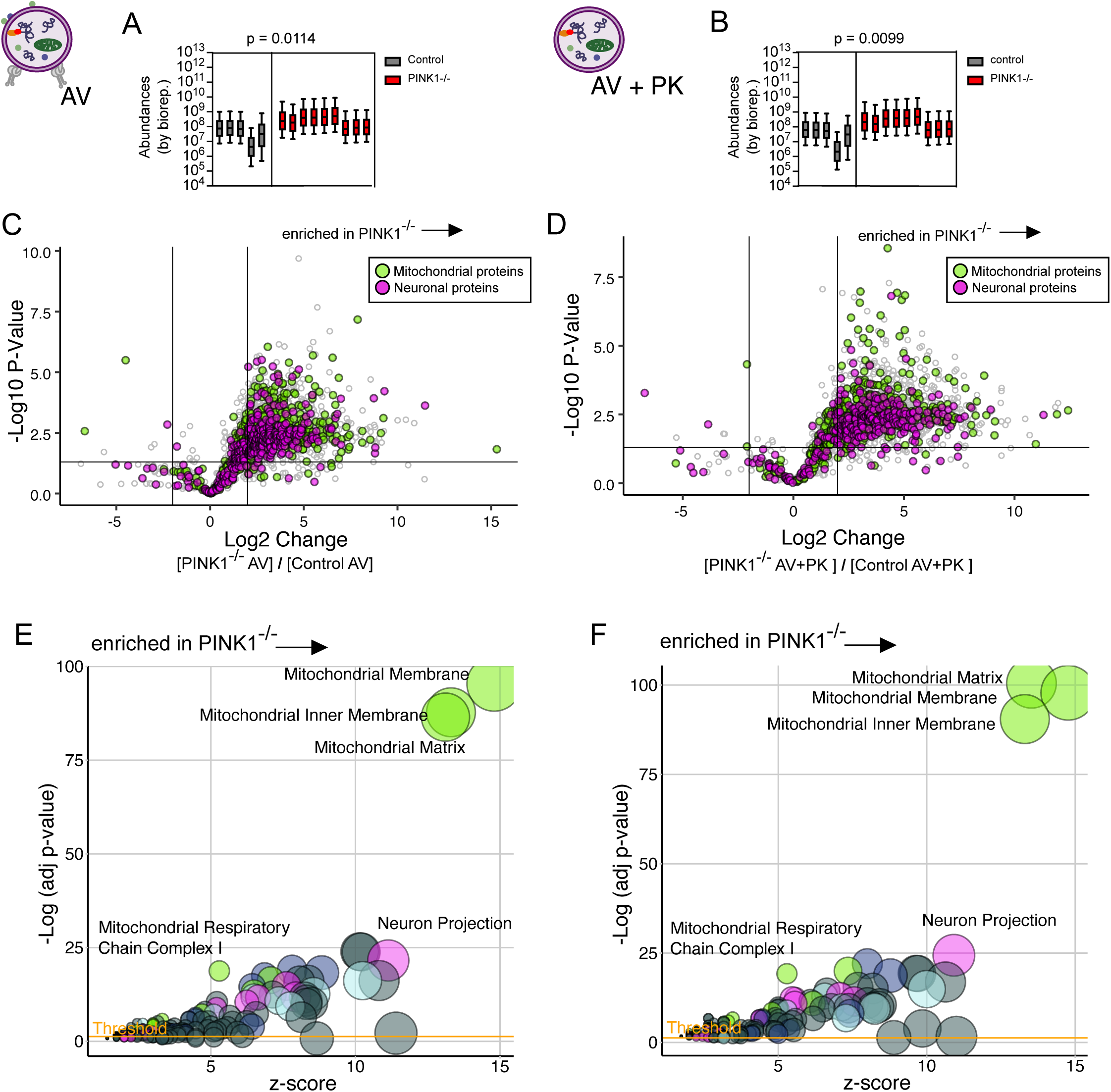

**Supplementary Figure 2.**
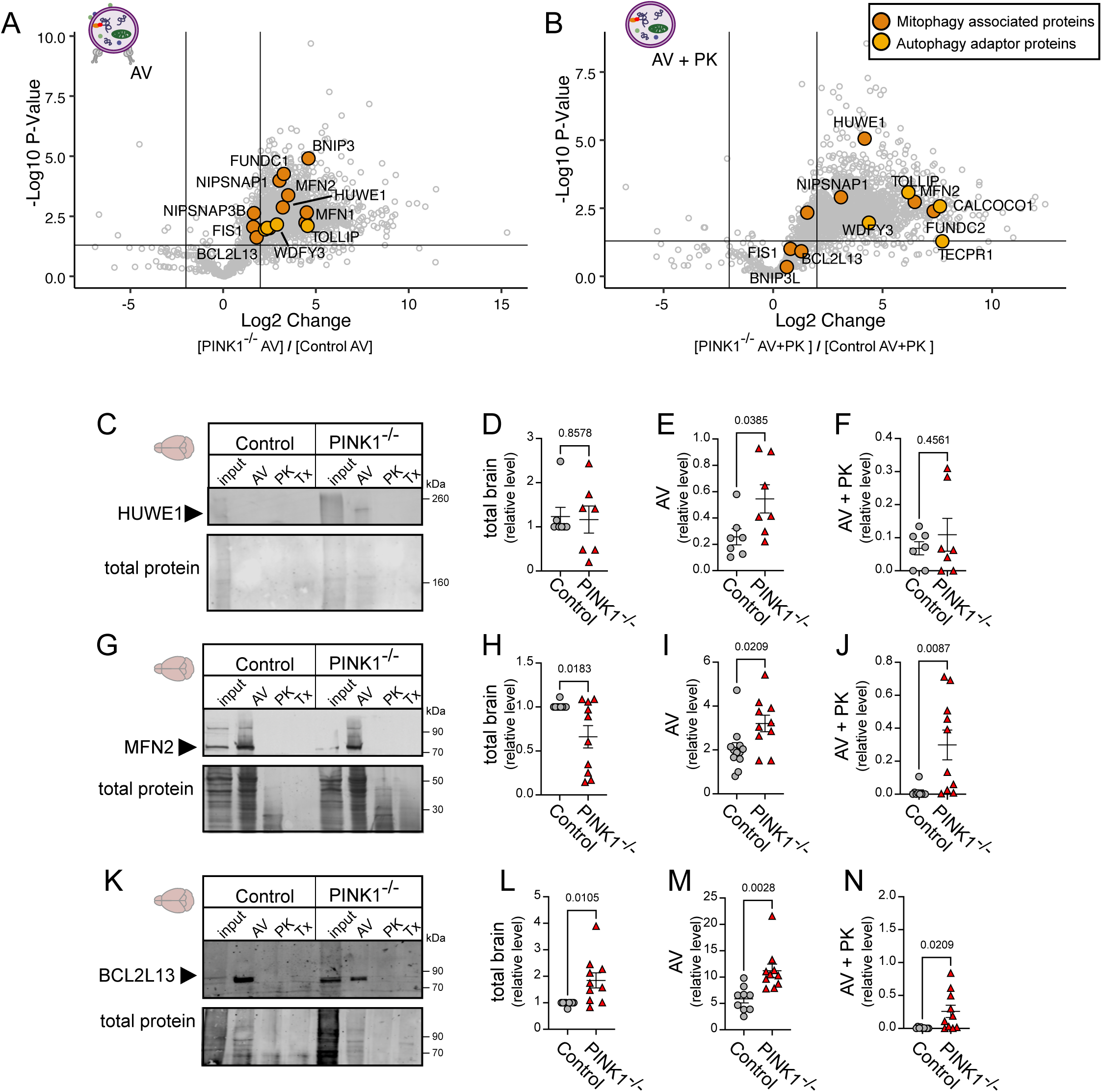

**Supplementary Figure 3.**
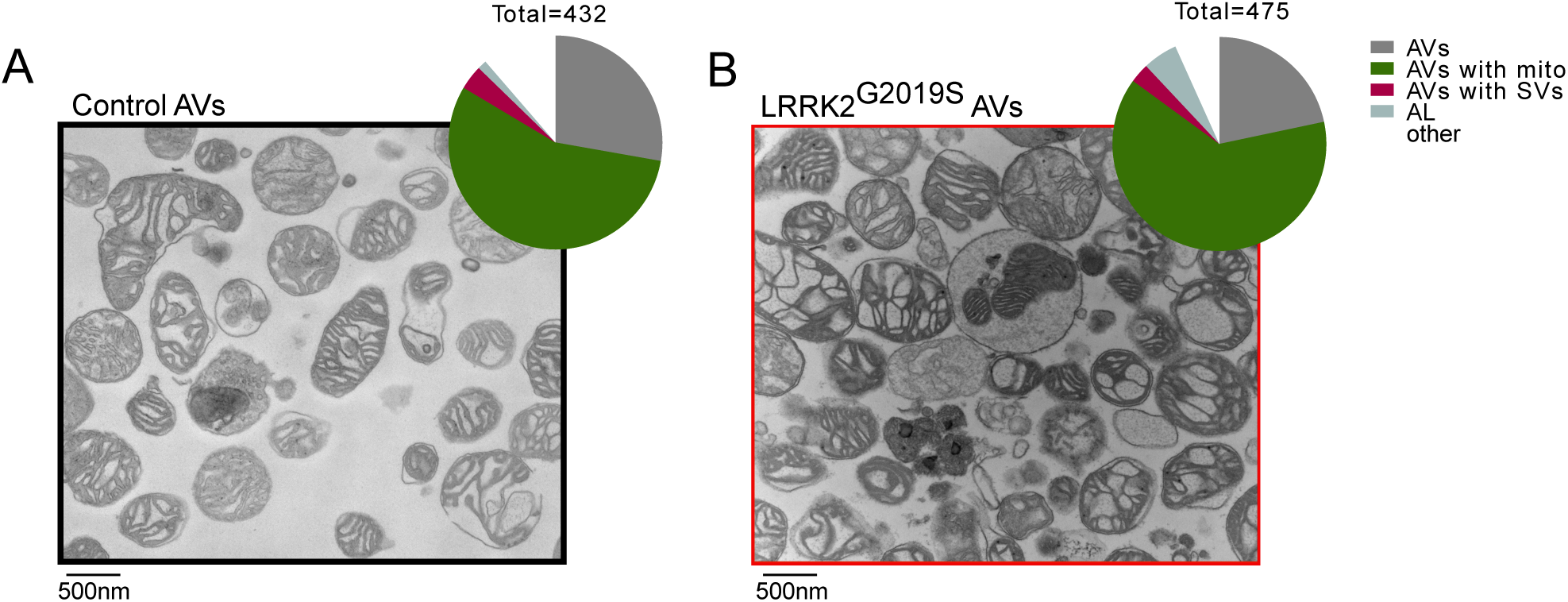

**Supplementary Figure 4.**
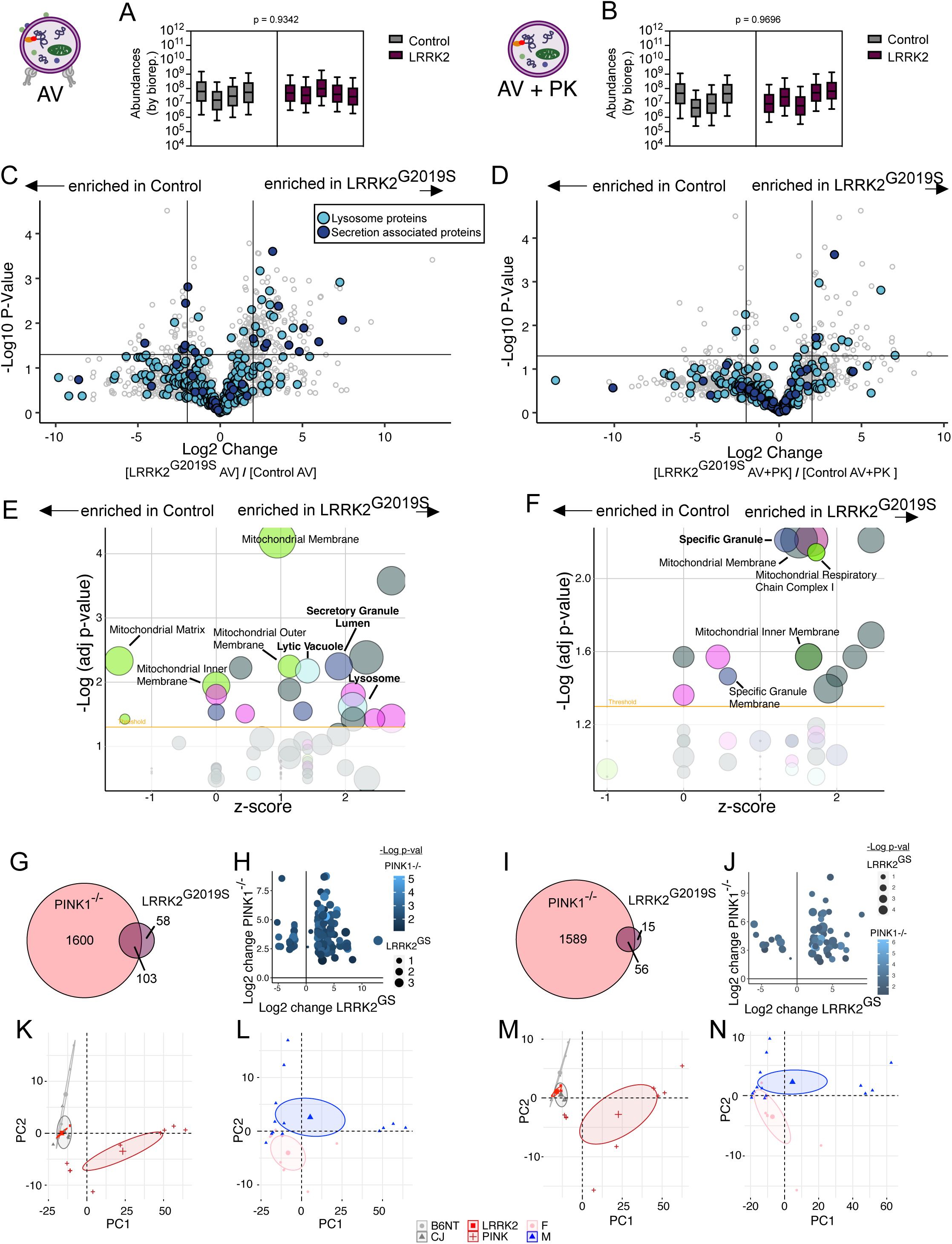

**Supplementary Figure 5.**
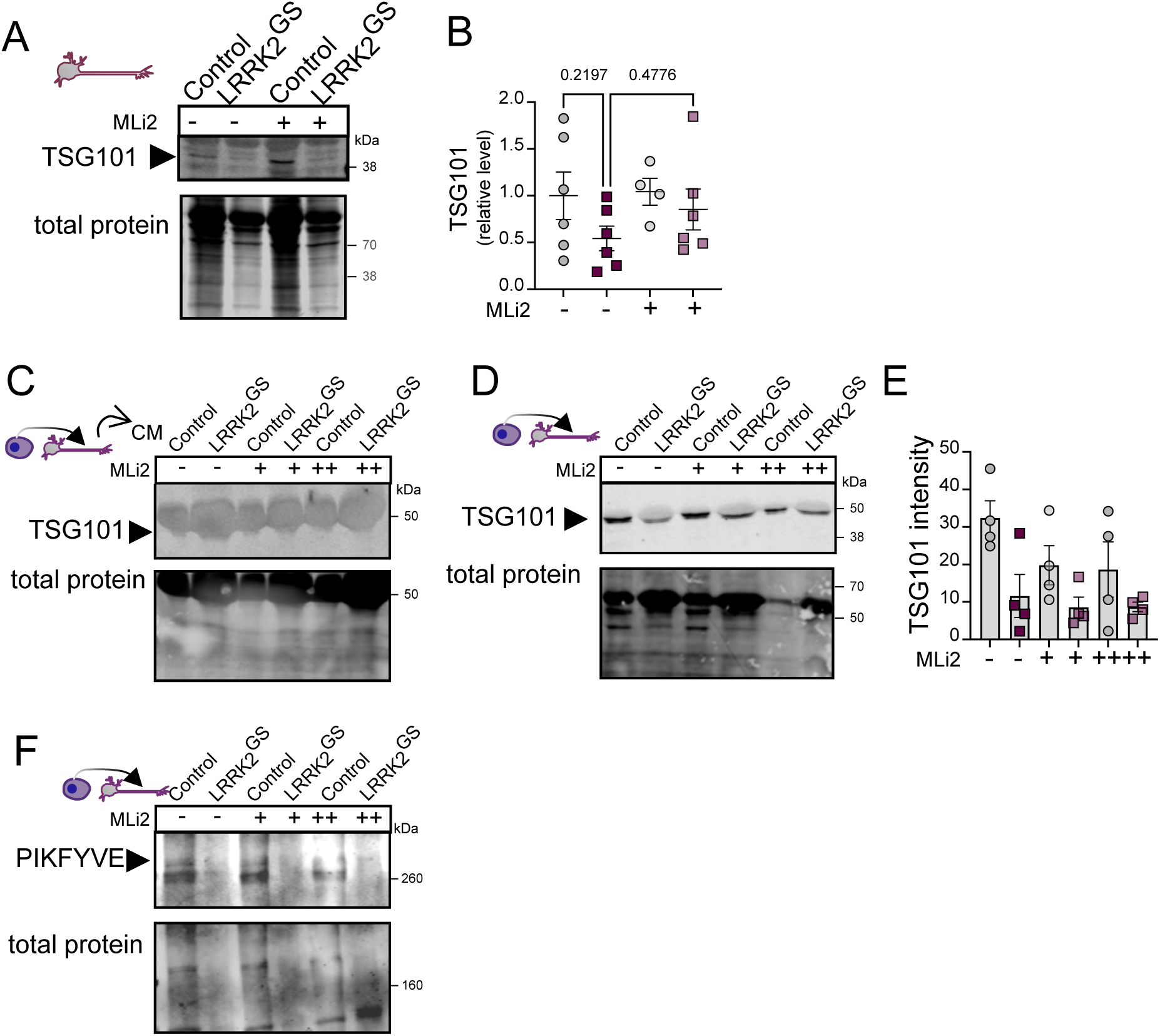

**Supplementary Figure 6.**
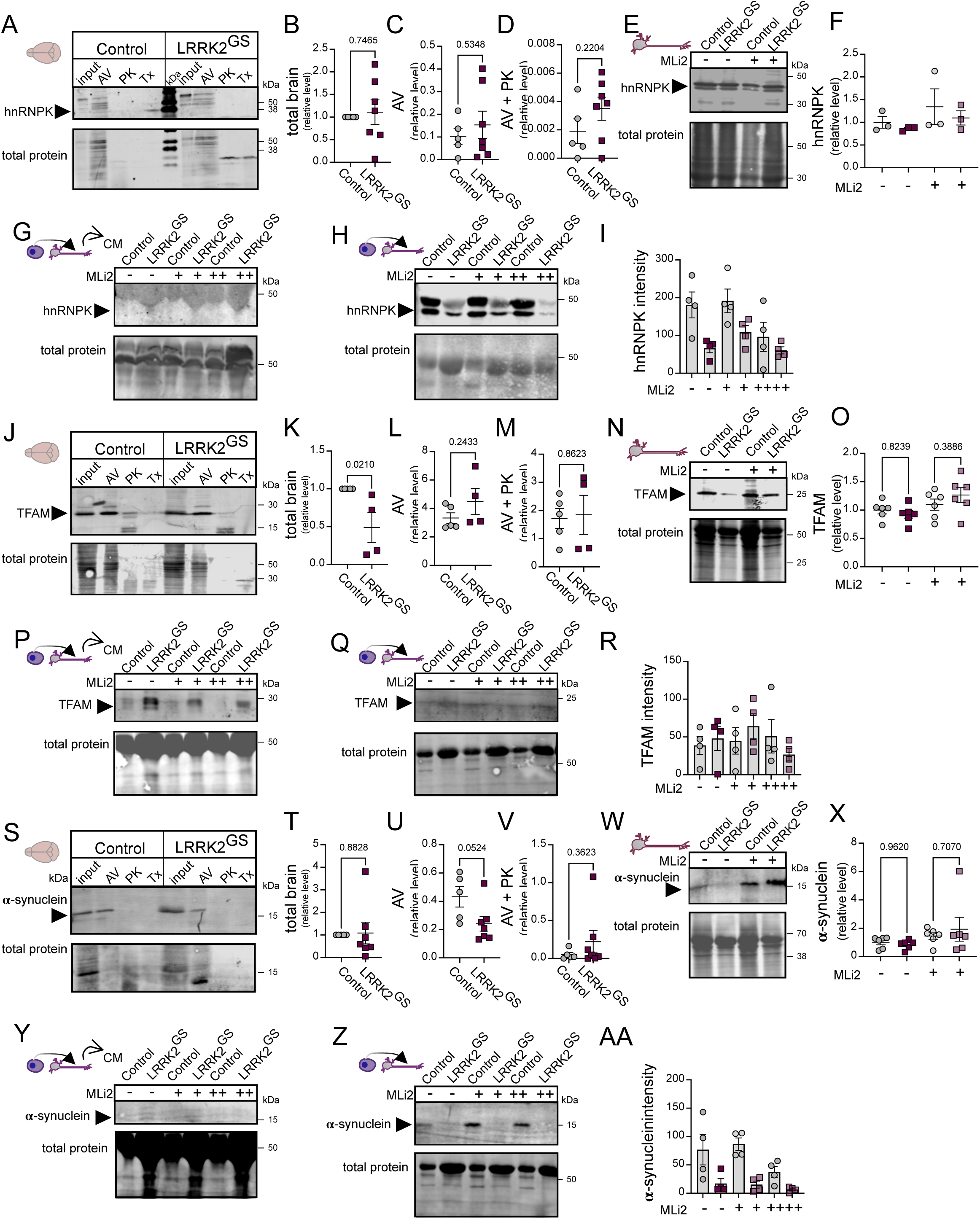

## References

1. Stavoe AKH, Holzbaur ELF. Autophagy in Neurons. Annu Rev Cell Dev Biol 2019; 35:477–500.

2. Wong E, Cuervo AM. Autophagy gone awry in neurodegenerative diseases. Nat Neurosci 2010; 13:805–11.

3. Fleming A, Rubinsztein DC. Autophagy in Neuronal Development and Plasticity. Trends Neurosci 2020; 43:767–79.

4. Kirkin V, Rogov V V. A Diversity of Selective Autophagy Receptors Determines the Specificity of the Autophagy Pathway. Mol Cell 2019; 76:268–85.

5. Lazarou M, Sliter DA, Kane LA, Sarraf SA, Wang C, Burman JL, Sideris DP, Fogel AI, Youle RJ. The ubiquitin kinase PINK1 recruits autophagy receptors to induce mitophagy. Nature 2015; 524:309–14.

6. Wong YC, Holzbaur ELF. Optineurin is an autophagy receptor for damaged mitochondria in parkin-mediated mitophagy that is disrupted by an ALS-linked mutation. Proc Natl Acad Sci U S A 2014; 111:E4439–48.

7. Ordureau A, Sarraf SA, Duda DM, Heo J-M, Jedrychowski MP, Sviderskiy VO, Olszewski JL, Koerber JT, Xie T, Beausoleil SA, et al. Quantitative Proteomics Reveal a Feedforward Model for Mitochondrial PARKIN Translocation and Ubiquitin Chain Synthesis. Mol Cell 2014; 56:462.

8. Heo J-M, Ordureau A, Paulo JA, Rinehart J, Harper JW. The PINK1-PARKIN mitochondrial ubiquitylation pathway drives a program of TBK1 activation and recruitment of OPTN and NDP52 to promote mitophagy Jin-Mi. Mol Cell 2015; 60:7–20.

9. Hara T, Nakamura K, Matsui M, Yamamoto A, Nakahara Y, Suzuki-Migishima R, Yokoyama M, Mishima K, Saito I, Okano H, et al. Suppression of basal autophagy in neural cells causes neurodegenerative disease in mice. Nature 2006; 441:885–9.

10. McWilliams TG, Prescott AR, Montava-Garriga L, Ball G, Singh F, Barini E, Muqit MMK, Brooks SP, Ganley IG. Basal Mitophagy Occurs Independently of PINK1 in Mouse Tissues of High Metabolic Demand. Cell Metab 2018; 27:439–449.e5.

11. Goldsmith J, Ordureau A, Harper JW, Holzbaur ELF. Brain-derived autophagosome profiling reveals the engulfment of nucleoid-enriched mitochondrial fragments by basal autophagy in neurons. Neuron 2022; 110:1–10.

12. Maday S, Wallace KE, Holzbaur ELF. Autophagosomes initiate distally and mature during transport toward the cell soma in primary neurons. J Cell Biol 2012; 196:407–17.

13. Ikenaka K, Kawai K, Katsuno M, Huang Z, Jiang Y, Iguchi Y. dnc-1 / dynactin 1 Knockdown Disrupts Transport of Autophagosomes and Induces Motor Neuron Degeneration. 2013; 8.

14. Katsumata K, Nishiyama J, Inoue T, Mizushima N, Katsumata K, Nishiyama J, Inoue T, Mizushima N, Takeda J, Yuzaki M. transport of autophagosomes in neuronal axons Dynein- and activity-dependent retrograde transport of autophagosomes in neuronal axons. 2010; 8627.

15. Neisch AL, Neufeld TP, Hays TS. A STR IPAK complex mediates axonal transport of autophagosomes and dense core vesicles through PP2A regulation. 2014; 216:441–61.

16. Cason SE, Carman P, Van Duyne C, Goldsmith J, Dominguez R, Holzbaur ELF. Sequential dynein effectors regulate axonal autophagosome motility in a maturation-dependent pathway. J Cell Biol 2021; 220:1–22.

17. Maday S, Holzbaur ELF. Autophagosome biogenesis in primary neurons follows an ordered and spatially regulated pathway. Dev Cell 2014; 30:71–85.

18. Anglade P, Vyas S, Javoy-Agid F, Herrero M, Michel P, Marquez J, Mouatt-Prigent A, Ruberg M, Hirsch E, Agid Y. Apoptosis and autophagy in nigral neurons of patients with Parkinson’s disease. Histol Histopathol 1997; 12:25–31.

19. Webb JL, Ravikumar B, Atkins J, Skepper JN, Rubinsztein DC. α-synuclein Is Degraded by Both Autophagy and the Proteasome. J Biol Chem 2003; 278:25009–13.

20. Sato S, Uchihara T, Fukuda T, Noda S, Kondo H, Saiki S, Komatsu M, Uchiyama Y, Tanaka K, Hattori N. Loss of autophagy in dopaminergic neurons causes Lewy pathology and motor dysfunction in aged mice. Sci Rep 2018; 8:1–10.

21. Di Maio R, Hoffman EK, Rocha EM, Keeny MT, Sanders LH, De Miranda BR, Zharikov A, Van Laar A, Stepan A, Lanz TA, et al. A central role for LRRK2 in idiopathic Parkinson Disease. Sci Transl Med 2018; 451.

22. Steger M, Diez F, Dhekne HS, Lis P, Nirujogi RS, Karayel O, Tonelli F, Martinez TN, Lorentzen E, Pfeffer SR, et al. Systematic proteomic analysis of LRRK2-mediated rab GTPase phosphorylation establishes a connection to ciliogenesis. Elife 2017; 6:1–22.

23. Taylor M, Alessi DR. Advances in elucidating the function of leucine-rich repeat protein kinase-2 in normal cells and Parkinson’s disease. Curr Opin Cell Biol 2020; 63:102–13.

24. Homma Y, Hiragi S, Fukuda M. Rab family of small GTPases: an updated view on their regulation and functions. FEBS J 2021; 288:36–55.

25. Boecker CA, Goldsmith J, Dou D, Cajka GG, Holzbaur ELF. Increased LRRK2 kinase activity alters neuronal autophagy by disrupting the axonal transport of autophagosomes. Curr Biol 2021; 31:2140–54.

26. Dou D, Smith EM, Evans CS, Boecker CA, Holzbaur ELF. Regulatory imbalance between LRRK2 kinase, PPM1H phosphatase, and ARF6 GTPase disrupts the axonal transport of autophagosomes Regulatory imbalance between LRRK2 kinase, PPM1H phosphatase, and ARF6 GTPase disrupts the axonal transport of autophagosomes. CellReports 2023; 42:112448.

27. Vincow ES, Thomas RE, Merrihew GE, Shulman NJ, Bammler TK, MacDonald JW, MacCoss MJ, Pallanck LJ. Autophagy accounts for approximately one-third of mitochondrial protein turnover and is protein selective. Autophagy 2019; 15:1592–605.

28. Di Rita A, Peschiaroli A, D′Acunzo P, Strobbe D, Hu Z, Gruber J, Nygaard M, Lambrughi M, Melino G, Papaleo E, et al. HUWE1 E3 ligase promotes PINK1/PARKIN-independent mitophagy by regulating AMBRA1 activation via IKKα. Nat Commun 2018; 9.

29. Otsu K, Murakawa T, Yamaguchi O. BCL2L13 is a mammalian homolog of the yeast mitophagy receptor Atg32. Autophagy 2015; 11:1932–3.

30. Kleele T, Rey T, Winter J, Zaganelli S, Mahecic D, Lambert HP, Ruberto F, Nemir M, Wai T, Pedrazzini T, et al. Distinct molecular signatures of fission predict mitochondrial degradation or proliferation. bioRxiv 2020; 2019.

31. Kestenbaum, M., Alcalay, R.N. (2017). Clinical Features of *LRRK2* Carriers with Parkinson’s Disease. In: Rideout, H. (eds) Leucine-Rich Repeat Kinase 2 (LRRK2). Advances in Neurobiology, vol 14. Springer, Cham. 10.1007/978-3-319-49969-7_2

32. Dou D, Smith EM, Evans CS, Boecker CA, Holzbaur ELF. Regulatory imbalance between LRRK2 kinase, PPM1H phosphatase, and ARF6 GTPase disrupts the axonal transport of autophagosomes. Cell Rep 2023; 42.

33. Solvik TA, Nguyen TA, Lin YHT, Marsh T, Huang EJ, Wiita AP, Debnath J, Leidal AM. Secretory autophagy maintains proteostasis upon lysosome inhibition. J Cell Biol 2022; 221:1–19.

34. Liang W, Sagar S, Ravindran R, Najor RH, Quiles JM, Chi L, Diao RY, Woodall BP, Leon LJ, Zumaya E, et al. Mitochondria are secreted in extracellular vesicles when lysosomal function is impaired. Nat Commun 2023; :1–18.

35. Leidal AM, Huang HH, Marsh T, Solvik T, Zhang D, Ye J, Kai FB, Goldsmith J, Liu JY, Huang Y-H, et al. The LC3-conjugation machinery specifies the loading of RNA-binding proteins into extracellular vesicles. Nat Cell Biol 2020; 22.

36. Zhong J, Ren X, Liu W, Wang S, Lv Y, Nie L, Lin R, Tian X, Yang X, Zhu F, et al. Discovery of Novel Markers for Identifying Cognitive Decline Using Neuron-Derived Exosomes. Front Aging Neurosci 2021; 13:1–14.

37. Sønder SL, Boye TL, Tölle R, Dengjel J, Maeda K, Jäättelä M, Simonsen AC, Jaiswal JK, Nylandsted J. Annexin A7 is required for ESCRT III-mediated plasma membrane repair. Sci Rep 2019; 9:1–12.

38. Li H, Liu N, Wang S, Wang L, Zhao J, Su L, Zhang Y, Zhang S, Xu Z, Zhao B, et al. Identification of a small molecule targeting annexin A7. Biochim Biophys Acta - Mol Cell Res 2013; 1833:2092–9.

39. Kowal J, Tkach M, Théry C. Biogenesis and secretion of exosomes. Curr Opin Cell Biol 2014; 29:116–25.

40. Assia S. PIKfyve: partners, significance, debates and paradoxes. Cell Biol Int 2008; 32(6):591–604.

41. de Lartigue J, Polson H, Feldman M, Shokat K, Tooze SA, Urbé S, Clague MJ. PIKfyve regulation of endosome-linked pathways. Traffic 2009; 10:883–93.

42. Sharma G, Guardia CM, Roy A, Vassilev A, Saric A, Griner LN, Marugan J, Ferrer M, Bonifacino JS, DePamphilis ML. A family of PIKFYVE inhibitors with therapeutic potential against autophagy-dependent cancer cells disrupt multiple events in lysosome homeostasis. Autophagy 2019; 15:1694–718.

43. Hessvik NP, Øverbye A, Brech A, Torgersen ML, Jakobsen IS, Sandvig K, Llorente A. PIKfyve inhibition increases exosome release and induces secretory autophagy. Cell Mol Life Sci 2016; 73:4717–37.

44. Osborne SL, Wen PJ, Boucheron C, Nguyen HN, Hayakawa M, Kaizawa H, Parker PJ, Vitale N, Meunier FA. PIKfyve negatively regulates exocytosis in neurosecretory cells. J Biol Chem 2008; 283:2804–13.

45. Leidal AM, Huang HH, Marsh T, Solvik T, Zhang D, Ye J, Kai F, Goldsmith J, Liu JY, Huang Y-H, et al. The LC3-conjugation machinery specifies the loading of RNA-binding proteins into extracellular vesicles. Nat Cell Biol 2020

46. Friedman LG, Lachenmayer ML, Wang J, He L, Poulose SM, Komatsu M, Holstein GR, Yue Z. Disrupted autophagy leads to dopaminergic axon and dendrite degeneration and promotes presynaptic accumulation of alpha-synuclein and LRRK2 in the brain. J Neurosci 2012; 32:7585–93.

47. Spillantini MG, Schmidt ML, Lee VMY, Trojanowski JQ, Jakes R, Goedert M. a - Synuclein in Lewy bodies. Nature 1997; 388:839–40.

48. Desplats P, Lee HJ, Bae EJ, Patrick C, Rockenstein E, Crews L, Spencer B, Masliah E, Lee SJ. Inclusion formation and neuronal cell death through neuron-to-neuron transmission of α-synuclein. Proc Natl Acad Sci U S A 2009; 106:13010–5.

49. Emmanouilidou E, Melachroinou K, Roumeliotis T, Garbis SD, Ntzouni M, Margaritis LH, Stefanis L, Vekrellis K. Cell-produced α-synuclein is secreted in a calcium-dependent manner by exosomes and impacts neuronal survival. J Neurosci 2010; 30:6838–51.

50. Nonaka T, Watanabe ST, Iwatsubo T, Hasegawa M. Seeded aggregation and toxicity of α-synuclein and tau: Cellular models of neurodegenerative diseases. J Biol Chem 2010; 285:34885–98.

51. Volpicelli-Daley L a, Luk KC, Patel TP, Tanik S a, Riddle DM, Stieber A, Meaney DF, Trojanowski JQ, Lee VM-Y. Exogenous α-synuclein fibrils induce Lewy body pathology leading to synaptic dysfunction and neuron death. Neuron 2011; 72:57–71.

52. Mougenot AL, Nicot S, Bencsik A, Morignat E, Verchère J, Lakhdar L, Legastelois S, Baron T. Prion-like acceleration of a synucleinopathy in a transgenic mouse model. Neurobiol Aging 2012; 33:2225–8.

53. Luk KC, Kehm V, Carroll J, Zhang B, O’Brien P, Trojanowski JQ, Lee VM-Y. Pathological α-Synuclein Transmission Initiates Parkinson-like Neurodegeneration in Non-transgenic Mice. Science 2012; 338:949–53.

54. Masuda-Suzukake M, Nonaka T, Hosokawa M, Oikawa T, Arai T, Akiyama H, Mann DMA, Hasegawa M. Prion-like spreading of pathological α-synuclein in brain. Brain 2013; 136:1128–38.

55. Ordureau A, Kraus F, Zhang J, An H, Park S, Ahfeldt T, Paulo JA, Harper JW. Temporal proteomics during neurogenesis reveals large-scale proteome and organelle remodeling via selective autophagy. Mol Cell 2021; 81.

56. Hoyer MJ, Smith IR, Paoli JC, Jiang Y, Paulo JA, Harper JW. Combinatorial selective ER-phagy remodels the ER during neurogenesis. BioRxiv 2023; June 26:1–39.

57. Behrends C, Sowa ME, Gygi SP, Harper JW. Network organization of the human autophagy system. Nature 2010; 466:68–76.

58. Gautier CA, Kitada T, Shen J. Loss of PINK1 causes mitochondrial functional defects and increased sensitivity to oxidative stress. Proc Natl Acad Sci U S A 2008; 105:11364–9.

59. Gispert S, Ricciardi F, Kurz A, Azizov M, Hoepken HH, Becker D, Voos W, Leuner K, Müller WE, Kudin AP, et al. Parkinson phenotype in aged PINK1-deficient mice is accompanied by progressive mitochondrial dysfunction in absence of neurodegeneration. PLoS One 2009; 4.

60. Shelke GV, Williamson CD, Jarnik M, Bonifacino JS. Inhibition of endolysosome fusion increases exosome secretion. J Cell Biol 2023; 222.

61. Hung S-T, Linares GR, Chang W-H, Eoh Y, Krishnan G, Mendonca S, Hong S, Shi Y, Santana M, Kueth C, et al. PIKFYVE inhibition mitigates disease in models of diverse forms of ALS Cell 2023; 186:786–802.e28.

62. Fussi N, Höllerhage M, Chakroun T, Nykänen NP, Rösler TW, Koeglsperger T, Wurst W, Behrends C, Höglinger GU. Exosomal secretion of α-synuclein as protective mechanism after upstream blockage of macroautophagy. Cell Death Dis 2018; 9.

63. Bayati A, Banks E, Han C, Luo W, Reintsch WE, Zorca CE, Shlaifer I, Del Cid Pellitero E, Vanderperre B, McBride HM, et al. Rapid macropinocytic transfer of α-synuclein to lysosomes. Cell Rep 2022; 40:111102.

64. Bae EJ, Kim DK, Kim C, Mante M, Adame A, Rockenstein E, Ulusoy A, Klinkenberg M, Jeong GR, Bae JR, et al. LRRK2 kinase regulates α-synuclein propagation via RAB35 phosphorylation. Nat Commun 2018; 9.

65. Schapansky J, Khasnavis S, DeAndrade MP, Nardozzi JD, Falkson SR, Boyd JD, Sanderson JB, Bartels T, Melrose HL, LaVoie M. Familial knockin mutation of LRRK2 causes lysosomal dysfunction and accumulation of endogenous insoluble α-synuclein in neurons Jason. Neurobiol Dis 2018; 111:26–35.

66. Ejlerskov P, Rasmussen I, Nielsen TT, Bergström AL, Tohyama Y, Jensen PH, Vilhardt F. Tubulin polymerization-promoting protein (TPPP/p25α) promotes unconventional secretion of α-synuclein through exophagy by impairing autophagosome-lysosome fusion. J Biol Chem 2013; 288:17313–35.

67. Larson ME, Greimel SJ, Amar F, LaCroix M, Boyle G, Sherman MA, Schley H, Miel C, Schneider JA, Kayed R, et al. Selective lowering of synapsins induced by oligomeric α-synuclein exacerbates memory deficits. Proc Natl Acad Sci U S A 2017; 114:E4648–57.

68. Kayed R, Dettmer U, Lesné SE. Soluble endogenous oligomeric α-synuclein species in neurodegenerative diseases: Expression, spreading, and cross-talk. J Parkinsons Dis 2020; 10:791–818.

69. Choi I, Zhang Y, Seegobin SP, Pruvost M, Wang Q, Purtell K, Zhang B, Yue Z. Microglia clear neuron-released α-synuclein via selective autophagy and prevent neurodegeneration. Nat Commun 2020; 11:1–14.

70. Pagano G, Ferrara N, Brooks DJ, Pavese N. Age at onset and Parkinson disease phenotype. Neurology 2016; 86:1400–7.

71. Bieri G, Brahic M, Bousset L, Couthouis J, Kramer NJ, Ma R, Nakayama L, Monbureau M, Defensor E, Schüle B, et al. LRRK2 modifies α-syn pathology and spread in mouse models and human neurons. Acta Neuropathol 2019; 137:961–80.

72. Volpicelli-Daley LA, Abdelmotilib H, Liu Z, Stoyka L, Daher JPL, Milnerwood AJ, Unni VK, Hirst WD, Yue Z, Zhao HT, et al. G2019S-LRRK2 expression augments α-synuclein sequestration into inclusions in neurons. J Neurosci 2016; 36:7415–27.

73. Fernandopulle MS, Prestil R, Grunseich C, Wang C, Gan L, Ward ME. Transcription-factor mediated differentiation of human iPSCs into neurons. Curr Protoc Cell Biol 2018; 79.

74. Schweppe DK, Prasad S, Belford MW, Navarrete-perea J, Bailey DJ, Huguet R, Jedrychowski MP, Rad R, Abbatiello SE, Woulters ER, et al. Characterization and optimization of multiplexed quantitative analyses using high-field asymmetric-waveform ion mobility mass spectrometry. Anal Chem 2019; 91:4010–6.

75. Tyanova S, Temu T, Sinitcyn P, Carlson A, Hein MY, Geiger T, Mann M, Cox J. The Perseus computational platform for comprehensive analysis of (prote)omics data. Nat Methods 2016; 13:731–40.

76. Calvo SE, Clauser KR, Mootha VK. MitoCarta2.0: An updated inventory of mammalian mitochondrial proteins. Nucleic Acids Res 2016; 44:D1251–7.

77. Xie Z, Bailey A, Kuleshov M V., Clarke DJB, Evangelista JE, Jenkins SL, Lachmann A, Wojciechowicz ML, Kropiwnicki E, Jagodnik KM, et al. Gene Set Knowledge Discovery with Enrichr. Curr Protoc 2021; 1:1–51.

78. Kuleshov M V., Jones MR, Rouillard AD, Fernandez NF, Duan Q, Wang Z, Koplev S, Jenkins SL, Jagodnik KM, Lachmann A, et al. Enrichr: a comprehensive gene set enrichment analysis web server 2016 update. Nucleic Acids Res 2016; 44:W90–7.

79. Chen EY, Tan CM, Kou Y, Duan Q, Wang Z, Meirelles G V., Clark NR, Ma’ayan A. Enrichr: Interactive and collaborative HTML5 gene list enrichment analysis tool. BMC Bioinformatics 2013; 14.

80. Koopmans F, van Nierop P, Andres-Alonso M, Byrnes A, Cijsouw T, Coba MP, Cornelisse LN, Farrell RJ, Goldschmidt HL, Howrigan DP, et al. SynGO: An Evidence-Based, Expert-Curated Knowledge Base for the Synapse. Neuron 2019; 103:217–234.e4

